# Nucleoredoxin regulates WNT signaling during pituitary stem cell differentiation

**DOI:** 10.1101/2025.01.30.635771

**Authors:** Michelle L. Brinkmeier, Leonard Y. M. Cheung, Sean P. O’Connell, Diana K. Gutierrez, Eve C. Rhoads, Sally A. Camper, Shannon W. Davis

**Affiliations:** Department of Human Genetics, University of Michigan, Ann Arbor, MI, 48109-5618, USA; Department of Biological Sciences, University of South Carolina, Columbia, SC, 29208, USA; Current address: Department of Physiology and Biophysics, Renaissance School of Medicine, State University of New York, Stonybrook, NY 11794, USA

**Keywords:** red-ox, WNT signaling, SOX2, growth hormone, cleft palate, Robinow syndrome, scRNAseq

## Abstract

Nucleoredoxin (*Nxn*) encodes a multi-functional enzyme with oxidoreductase activity that regulates many different signaling pathways and cellular processes in a redox-dependent manner. Rare *NXN* mutations are reported in individuals with recessive Robinow syndrome, which involves mesomelic skeletal dysplasia, short stature, craniofacial dysmorphisms, and incompletely penetrant heart and palate defects. Here we report that *Nxn* is expressed in the ventral diencephalon and developing pituitary gland, and that *Nxn* deficient mice have pituitary dysmorphology and craniofacial abnormalities that include defects in the skull base and cleft palate. *Nxn* mutant mice exhibit reduced WNT signaling and reduced differentiation of pituitary stem cells into hormone-producing cells. These results suggest patients with Robinow syndrome could benefit from evaluation by endocrinologists for pituitary structural imaging and hormone insufficiency.

**Highlights:**

*Nxn* deficiency causes neonatal lethality, cleft palate, craniofacial abnormalities, and other structural birth defects.
*Nxn* deficiency causes pituitary dysmorphology.
Analysis of scRNA seq reveals delayed trajectory during early pituitary cell differentiation.
*Nxn* deficiency reduces canonical and non-canonical Wnt signaling in developing pituitary glands.
*Nxn* deficiency reduces stem cell differentiation into pituitary hormone-producing cells.

## Introduction

### Nucleoredoxin function and human disease

Maintenance of reduction-oxidation equilibrium is important to protect cells from oxidant damage and to initiate damage repair ^1^. Nucleoredoxin (NXN) is a thioredoxin that can regulate cellular redox homeostasis ^2,3^. NXN interacts with multiple proteins to regulate several pathways. These include disheveled (DVL), protein phosphatase (PP2A), phosphofructokinase (PFK1), endoplasmic reticulum transport protein (SEC63), regulator of interleukin and Toll-like receptor signaling (MYD88), regulation of actin polymerization (FLII), and calcium, calmodulin dependent protein kinase (CAMK2A) ^4^. DVL integrates WNT signals through both the canonical WNT-β-catenin and non-canonical WNT-planar cell polarity (PCP) pathways. NXN acts as a redox sensor; in the presence of reactive oxygen species, its cysteine residues are reduced, which results in release of disheveled ^3^. This can result in reduced or increased WNT signaling^5,6^.

Biallelic mutations in *NXN* are a rare cause of Robinow Syndrome, which is characterized by short stature, skeletal dysplasia that includes mesomelic limb shortening, and mild facial dysmorphology ^7–9^. Some Robinow syndrome cases are associated with cleft lip and/or cleft palate ^10^. Robinow syndrome can also be caused by mutations in other genes that are important in the WNT/PCP signaling pathway, including *WNT5A,* the disheveled genes *DVL1, DVL2,* and *DVL3,* and the WNT co-receptors *FZD2* and *ROR2* ^8,9^. Like *NXN,* bi-allelic loss of function mutations in the *WNT5A* co-receptor *ROR2* causes Robinow syndrome, while mutations in the other causative genes are dominant. Carboxy-terminal, dominant negative mutations in *DVL* genes account for most cases to date (∼33%), and missense mutations in *WNT5A* and *FZD2* each account for about 9.5% of cases.

### Role of WNT signaling in pituitary, hypothalamic, and palate development

WNT signaling is required for many aspects of pituitary development: 1) the expansion of Rathke’s pouch, which is the primordium for the anterior (pars distalis) and intermediate lobes (pars intermedia) of the pituitary gland ^11–13^, 2) the development of the posterior lobe (pars nervosa) and pituitary stalk, which is surrounded by the pars tuberalis ^14,15^, 3) postnatal expansion of the pituitary gland ^16^, and 4) pituitary remodeling in mature animals ^17,18^. It is also important for patterning and differentiation in the hypothalamus, especially the neurons that regulate anterior pituitary function ^19^. Finally, both canonical and noncanonical WNT signaling are necessary for normal palate development ^6^. Activation of WNT signaling in pituitary stem cells during development can cause adamantinomatous craniopharyngioma in mice and humans ^20,21^. WNT signaling may contribute to hypothalamic hamartoma, early onset, non-progressive pituitary adenoma, and both functioning and non-functioning symptomatic pituitary adenomas ^22–25^.

The mechanisms whereby WNT signaling affects pituitary growth in normal and disease states are beginning to emerge. Early studies demonstrated roles for *Wnt4, Wnt5a,* and *Wnt11* ^13,26,27^. Pinpointing the role of individual WNT genes in pituitary development has been challenging because there are 19 WNT genes in mammals, and at least 14 of them are expressed in pituitary gland ^16,28,29^. However, recent studies have elegantly demonstrated that pituitary stem cells expressing SOX2 secrete WNTs to stimulate nearby lineage-committed progenitors to proliferate during the postnatal period of pituitary growth ^16^.

Many transcription factors that are critical for hypothalamic and/or pituitary development are regulated by β-catenin, and disruption of these genes can affect pituitary development and/or function, including TCF7L1, TCF7L2, PROP1, POU1F1, PITX2, LEF1, NR5A1, FOXO1, and FOXO3 ^29–41^. In addition, SOX2 and SOX3 can suppress WNT signaling ^42^, and mutations in these genes also cause hypopituitarism ^24,43^.

### Nucleoredoxin and hypopituitarism

Mutations in approximately 90 genes can cause hypopituitarism ^44,45^, but many cases do not yet have mutations in known genes. We identified 50 genes that cause pituitary developmental abnormalities in embryonic lethal or sub-viable mouse knockouts that have not previously been associated with pituitary abnormalities ^44^. We hypothesized that some of these genes could be associated with syndromic hypopituitarism in humans. We discovered that mice homozygous for nucleoredoxin (*Nxn*) loss-of-function have anterior pituitary dysmorphology, and we show here that the differentiation of stem cells to hormone producing cells is reduced. These results are important because they reveal a role for NXN in pituitary development and suggest that individuals with biallelic *NXN* mutations may benefit from endocrine evaluation.

## Materials and methods

### Mice

All procedures using mice were approved by the University of Michigan Committee on Use and Care of Animals, and all experiments were conducted in accordance with the principles and procedures outlined in the National Institutes of Health Guidelines of the Care and Use of Experimental Animals.

*Nxn^em1(IMPC)J^* mice were generated at the Jackson Lab by CRISPR/Cas gene editing in the C57BL/6N genetic background. They contain a 5 bp deletion of ATCGC in exon 2 beginning at Chromosome 11 negative strand position 76278553bp (GRCm38). This mutation is predicted to cause amino acid sequence changes after residue 133 and early truncation 21 amino acids later: p.Ser134fsAsnTer155. Sperm from this strain were purchased from Mutant Mouse Resource and Research Center at University of California, Davis (<u>RRID:MMRRC_048884-UCD</u>). In vitro fertilization was performed by The University of Michigan Transgenic Animal Model Core facility using eggs from female C57BL/6NJ donors (<u>RRID:IMSR_JAX:005304</u>). Founder mice were mated with C57BL/6J (<u>RRID:IMSR_JAX:000664</u>).

Mice were maintained at the University of Michigan through heterozygous crosses. All mice were housed in a 12-hour light, 12-hour dark cycle in ventilated cages with unlimited access to tap water and Purina 5020 chow. The morning after conception is designated e0.5 and the day of birth is designated as P0.

### Genotyping

Genomic DNA was amplified by PCR in separate reactions for the wild type and mutant alleles. The forward oligonucleotides for the wild type (5’ AGTCTCCAACATTCCATCGC 3’) and mutant alleles (5’ ACCGAGTCTCCAACATTCCT 3’) were amplified with the reverse oligonucleotide primer (5’ AGTGTTGGTAGCTGGGTTCT 3’) under the following conditions: 92 C 2 min followed by 30 cycles of: 90 C 10 sec, 57 C 30 sec, 72 C 30 sec, and a final hold at 4 C. The wild type and mutant alleles produced 378 and 377 bp bands, respectively.

### Single cell RNA sequencing (scRNAseq)

We dissected pituitary glands from control (1 wild type, 1 heterozygote) and 2 *Nxn* mutants at e14.5 and dispersed the pooled tissue to single cells in 50µl of a solution containing 1mg/ml papain (Roche, cat# 10108014001), 5mM L-cysteine, 1mM EDTA, and 0.6mM 2-mercaptoethanol. Dissociation was performed at 37°C for 5 minutes, with titration every 2 minutes. The samples were neutralized in 75µl of Neurobasal Media (Invitrogen, cat# 12348017) with 10% fetal bovine serum (Corning, cat# MT35010CV). The single cells were treated with 0.75µl of RNAse free Recombinant DNase I (Roche, cat#4716728001) at 37°C for 5 minutes. Single cells were pelleted in a pre-chilled microfuge for 5 minutes at 300 RCF. The supernatant was aspirated, and the cells were resuspended in 50µl Neurobasal Media with 1% fetal bovine serum. Cell viability was analyzed using the Countess 3 cell counter (Invitrogen). Viability was excellent: 95% for control and 90% for mutant. Samples were processed to create cDNA libraries at the University of Michigan Advanced Genomics Core according to a standard pipeline ^46^ using the 10x Genomics Chromium platform and controller, following the manufacturer’s instructions for Single Cell 3’ v3 reagents, and sequenced on the NovaSeq6000 platform with an S4 chip. Sequencing reads were processed, and cells were clustered using Seurat 4.1.0. For controls and mutants, respectively, the number of cells sequenced was 19,625 and 15,918, the mean reads/cell was 27,387 and 34,783, and median genes/cell was 911 and 777. Single-cell RNA sequencing (scRNAseq) data described here are available on the National Center for Biotechnology Information Gene Expression Omnibus ^47^ using accession numbers GSM8628052 and GSE281783. The control pool was reported previously, GSM7864907 and GSE246211 ^44^.

### Immunohistochemistry and in situ hybridization

Immunostaining was carried out on tissues fixed in paraformaldehyde, embedded in paraffin, and sectioned and stained as previously described ^48^. Antibody sources, dilutions and detection methods are listed in **Supplementary Material, Table 1**.

RNAscope in situ hybridization for *Nxn* (Cat # 507321) was performed as previously described, including the RNAscope 2.5 HD Brown Detection kit, from Advanced Cell Diagnostics (Cat#322360).

### Skeletal staining

Skeletal preparations were stained with alcian blue and alizarin red to visualize cartilage and bone, respectively ^49^. The areas of cartilage and bone staining were calculated with Image J2, Fiji ^50^.

### Statistical Analysis

We used Prism-GraphPad to create graphs and compute statistics for cell number. Data are presented as standard error of the mean (SEM) and evaluated with Student’s unpaired t test or multiple unpaired t test.

## Results

### Nucleoredoxin mutations and pituitary dysmorphology

NXN encodes a 435 amino acid protein that is conserved in vertebrates and is 98% identical in mice (Chr 11) and humans (Chr 17). It contains a thioredoxin fold that catalyzes disulfide bond formation and isomerization (**Figure 1A**). Five loss-of-function variants have been reported in humans ^8,9,51^. Mouse *Nxn* knockouts and a hypomorphic (reduced function) allele have been reported, although the pituitary gland and hypothalamus have not been examined (**Supplemental Figure 1**) ^1,52–56^. We characterized the *Nxn^em1(IMPC)J^* allele, which we refer to here as *Nxn^-/-^.* It contains a 5 bp deletion, c.479del483 (NM_008750), which is predicted to produce a truncated protein, p.Ser134fsAsnTer155 that would lack functional domains. We intercrossed *Nxn^+/-^* mice and obtained a normal distribution of genotypes at e10.5-e11.5 (N=76, p=0.11), e14.5 (N=118, p=0.99), and e17.5-e18.5 (N=82, p=0.19), but no mutants at 2-3 wk (N=116, p = 0.001). (**Supplemental Table 2)**. Other *Nxn* loss of function alleles are lethal at birth, and a hypomorphic allele *Nxn^11Jus13^* exhibited reduced viability by weaning ^1,52,53^. The lethality has been attributed to heart and palate defects, while the placenta is normal.

**Figure 1.**
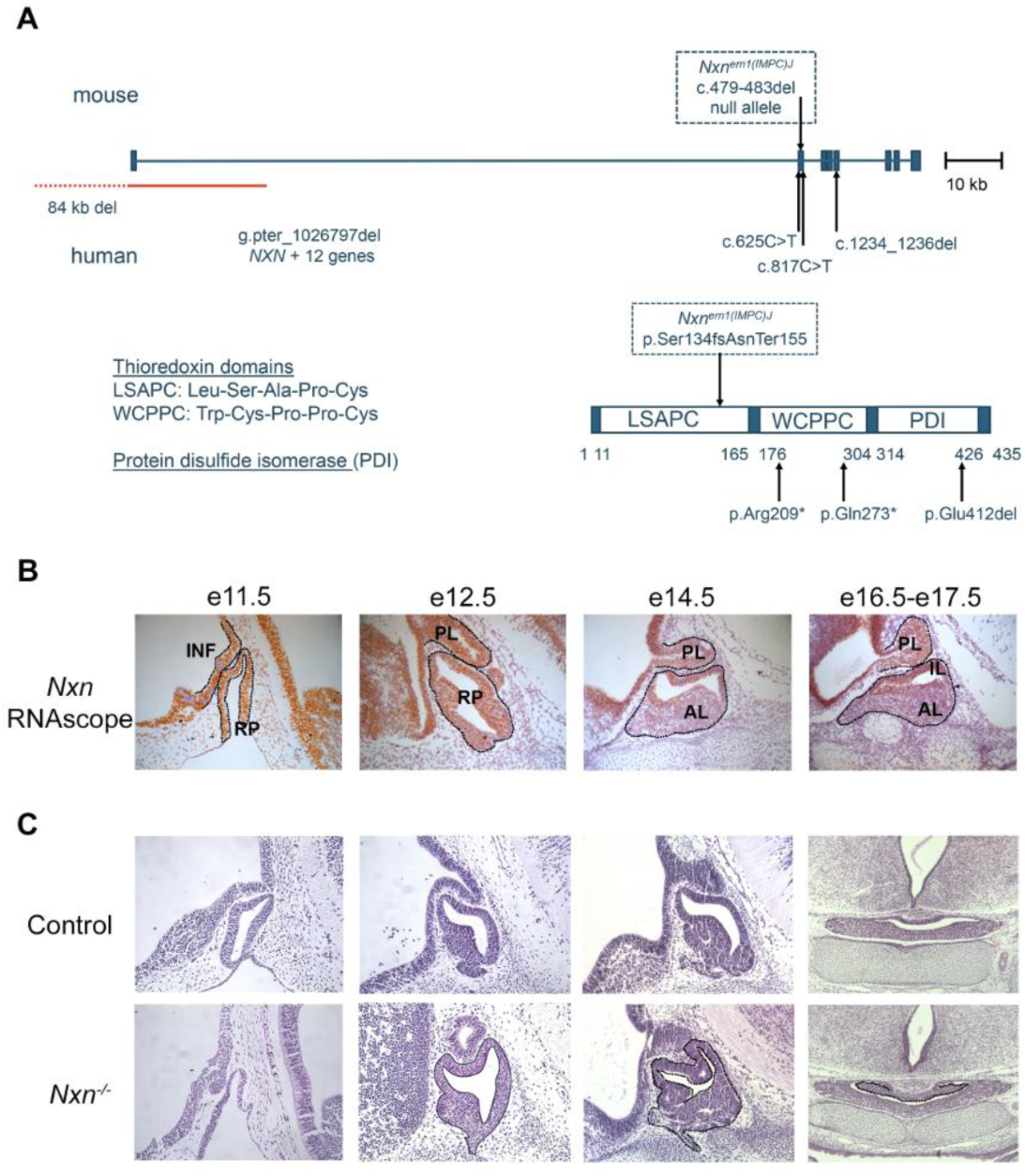
Nucleoredoxin loss of function mutations, pituitary gene expression, and dysmorphology. A. The mouse *Nxn* gene is shown with the positions of the variants in the *Nxn^em1/J^* (referred to here as *Nxn^-/-^*) allele. Clinically reported mutations in NXN are indicated below the diagram ^8,9,51^. A diagram of the protein domains shows the thioredoxin domains, LSAPC (Leu-Ser-Ala-Pro-Cys) and WCPPC (Trp-Cys-Prop-Pro-Cys), and the protein disulfide isomerase domain (PDI). B. Control and *Nxn^-/-^* fetuses were collected at e11.5, e12.5, e14.5 and e16.5-e17.5. *Nxn* expression was assessed with RNAscope in situ hybridization in control tissues (orange). Transcripts were detected in the infundibulum (INF), Rathke’s pouch (RP), the posterior lobe (PL), and the anterior and intermediate lobes (IL, AL respectively). C. Sections stained with eosin and hematoxolin revealed dysmorphology in Rathke’s pouch tissue (black outline) in mutants at e11.5, e12.5 and e14.5 (sagittal). Dysmorphology was evident in e16.5-e17.5 sections (coronal), (black outline).

*Nxn* is expressed broadly ^2^, and we detected it in cDNA libraries from pituitaries collected at e12.5 and e14.5 ^29^. We used RNAscope in situ hybridization to characterize the spatial and temporal expression of *Nxn* during pituitary and hypothalamic development (**Figure 1B**). Expression was detected in Rathke’s pouch, the precursor of the anterior and intermediate lobes of the pituitary gland, and in the ventral diencephalon and developing infundibulum at e11.5 and e12.5. At e14.5-e17.5 expression persisted in the ventral diencephalon and posterior lobe, but expression in the anterior lobe was strong in the marginal zone or stem cell niche, and weak in the cells beginning to differentiate (located more ventrally and rostrally than the stem cells).

Histological analysis of *Nxn^-/-^* embryos revealed dysmorphology of Rathke’s pouch from e11.5 through e17.5 (**Figure 1C**). At 11.5 the mutant pouch was thin and poorly developed. At e12.5 the dorsal aspect of the pouch was highly dysmorphic, suggesting over-proliferation of the stem cell niche and/or failure of progenitors to delaminate and move into the parenchyma of the developing anterior lobe. The dysmorphology we noted in mutants at e14.5 was like the mutant embryos presented in the Deciphering the Mechanisms of Developmental Disorders (DMDD) database using high-resolution episcopic microscopy (HREM) images ^44,57,58^. The developing posterior lobe appeared smaller than normal.

### Craniofacial abnormalities and fully penetrant cleft palate in *Nxn* mutants

We collected *Nxn ^-/-^* mutants at e18.5 and noted variable penetrance of facial clefting and micro-ophthalmia or anophthalmia (**Supplemental Fig. 2**). We visually inspected the palate and noted completely penetrant cleft palate in mutants (N=7/7; and 6/7 were severe). To quantify alterations in craniofacial structures we prepared skeletons from e18.5 embryos with alizarin red and alcian blue staining (**Fig. 2A-F**). No abnormalities were noted in wild type (N=4) or *Nxn^+/-^* heterozygous embryos (N=16), but the homozygous mutants (*Nxn ^-/-^*) had fully penetrant craniofacial defects and incomplete ossification of the basisphenoid bone within the skull base (100%, N=7). The length of the head, nasal bone, and mandible were all significantly shorter in mutants (**Fig. 2G)**. We quantified the relative size of the opening in the basisphenoid bone to the area of the entire bone, and we found that the size of the opening was variable in mutants, but on average it was ∼8x larger in mutants (5.6% ± 0.02) than wild type or heterozygotes (0.06 ± 0.01%).

**Figure 2.**
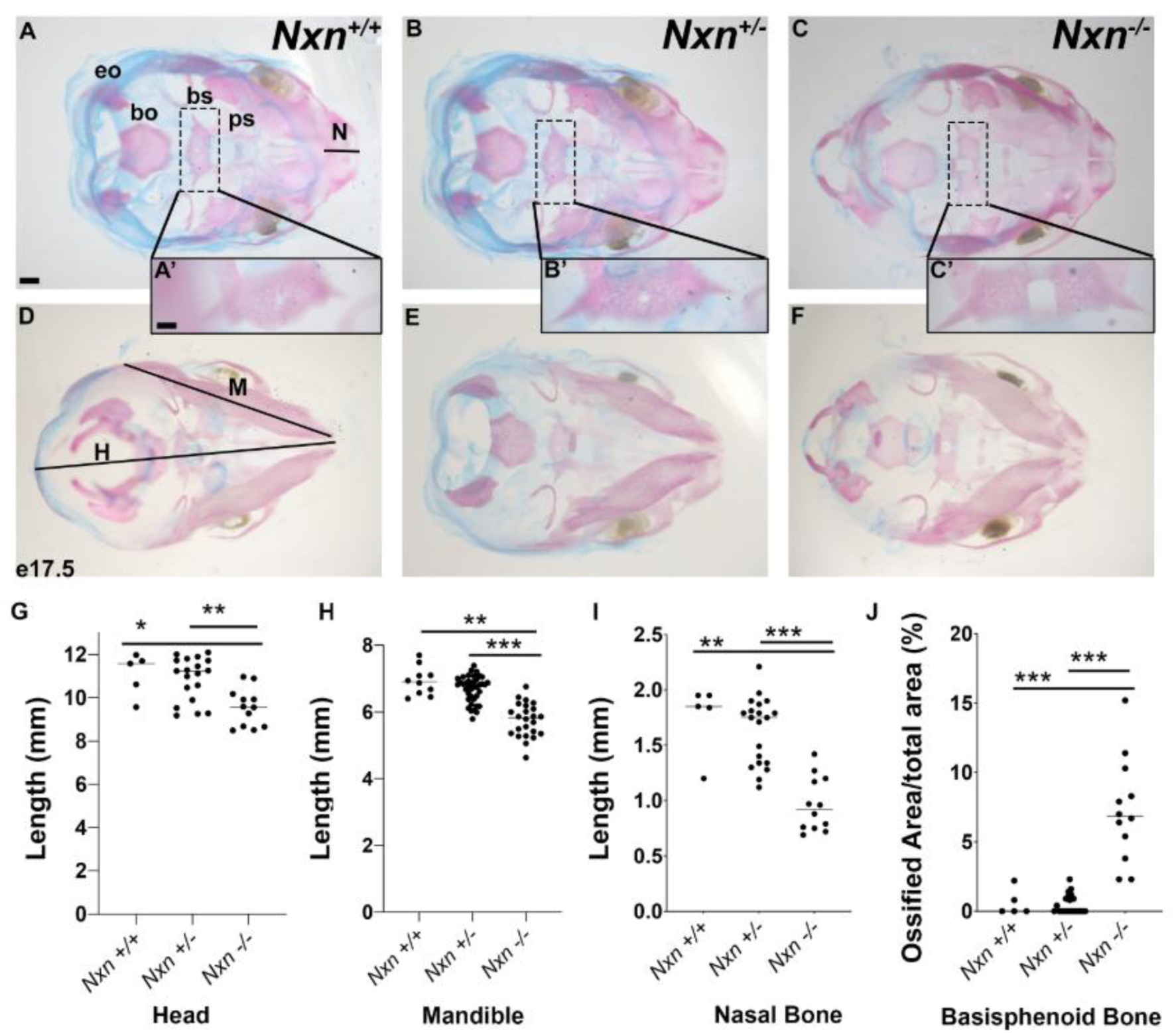
Nucleoredoxin deficiency causes craniofacial dysmorphology and incomplete ossification of the basisphenoid bone. Fetuses were collected at e18.5, and skeletons were prepared with alcian blue to visualize cartilage and alizarin red for bone. Preparations were photographed from dorsal (A, B, C) and ventral perspectives (D, E, F). The length of the nasal bone (N), head (H), and mandible (M) were quantified. The exoccipital (eo), basioccipital (bo), basisphenoid (bs), presphenoid (ps) bones are indicated, and the insets show an enlargement of the basisphenoid bone (A’, B’, C’). The basisphenoid bone was incompletely ossified in *Nxn^-/-^* embryos, and the size of the opening was measured relative to the total area of this element (G). Bone and opening measurements are indicated for fetuses collected at e18.5 of each genotype: wild type (N=4), heterozygote (N=16), and mutant (N=7). The p values are designated as *** = p<0.01, ** = p<0.05.

### scRNAseq analysis of gene expression

To assess the effect of *Nxn* deficiency on gene expression in the pituitary gland we conducted scRNAseq on control and mutant pools of cells from dissected pituitaries of e14.5 fetuses. 15 cell clusters were identified in controls and mutants (**Figure 3A**). We used our knowledge of cell-type specific markers to define the clusters (**Supplemental Fig. 3**) ^15,46,59^. In general, the pituitary cell clusters associated with Rathke’s pouch are defined by transcripts for *Epcam* (**Fig. 3B**) ^60^, and the transcription factors *Lhx3, Pitx1, Prop1, Pou1f1, Gata2* and *Tbx19* (**Fig. 3C**). The neural clusters are defined by transcripts for the transcription factors *Rax* and *Nkx2.1*, and the absence of *Epcam1* expression.

**Figure 3.**
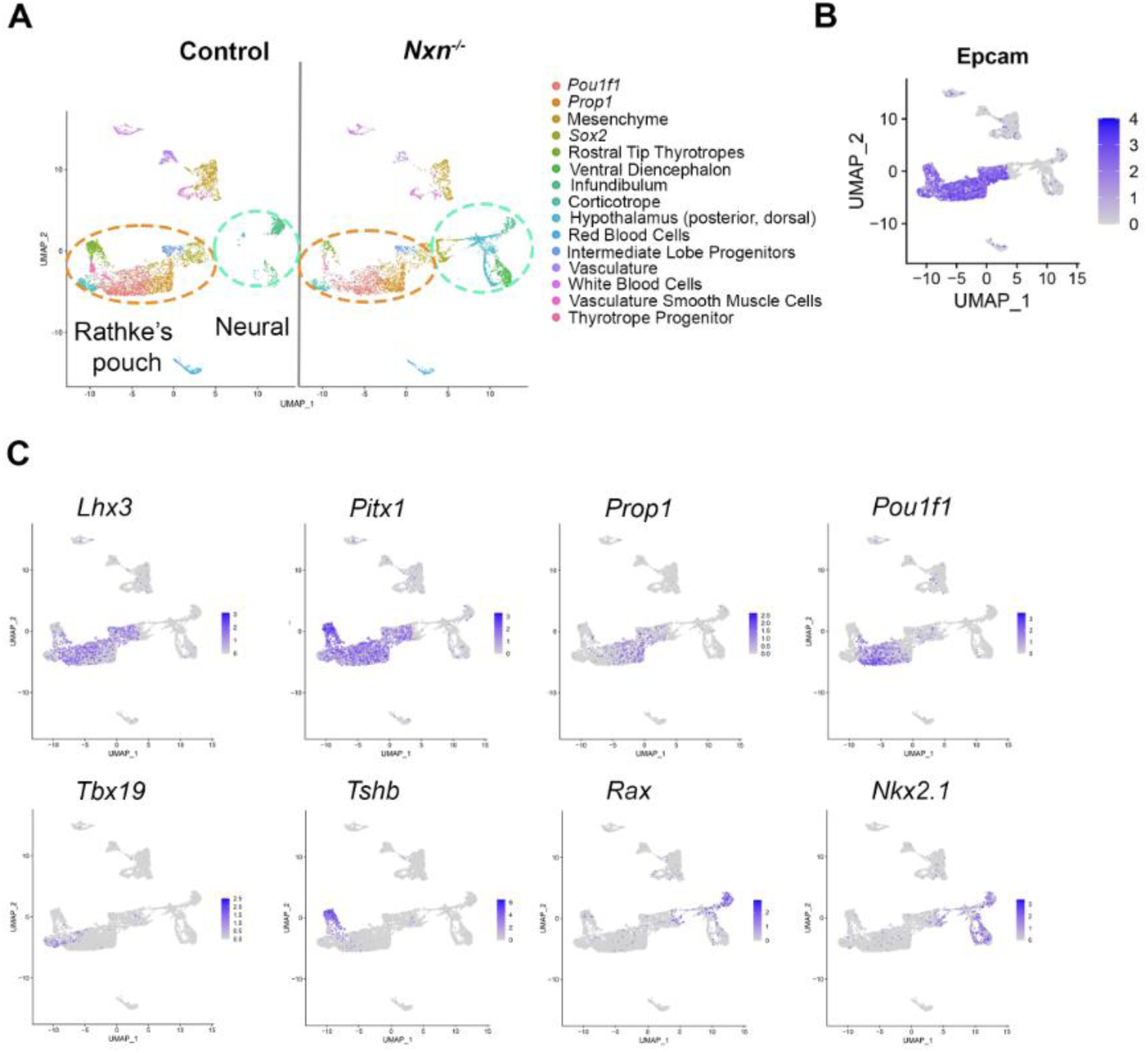
scRNAseq cell clusters are defined by *Epcam1,* various transcription factors and hormone gene expression. A. UMAP plots of scRNAseq data from pituitaries collected from control and *Nxn^-/-^* fetuses at e14.5. B. UMAP plot illustrating strong *Epcam* expression in Rathke’s pouch derived cell types. C. UMAP plots of lineage specific transcription factors that were used to assign cell cluster identity.

As expected, *Nxn* transcripts were broadly detected in controls, but not in mutants (**Fig. 4A**). The clusters with the highest levels of *Nxn* expression in Rathke’s pouch were *Sox2* expressing stem cells, *Prop1* expressing progenitors, and intermediate lobe progenitors. *Nxn* transcripts were also abundant in the ventral diencephalon, infundibulum, and surrounding mesenchyme. Direct effects of *Nxn* deficiency on gene expression would be expected in these cell types. Little or no *Nxn* transcripts were detected in the corticotropes, committed *Pou1f1* cells, or rostral tip thyrotropes (pars tuberalis), which are derived from Rathke’s pouch and surround the pituitary stalk ^15^. This is consistent with the results of the *Nxn* in situ hybridization.

**Fig. 4.**
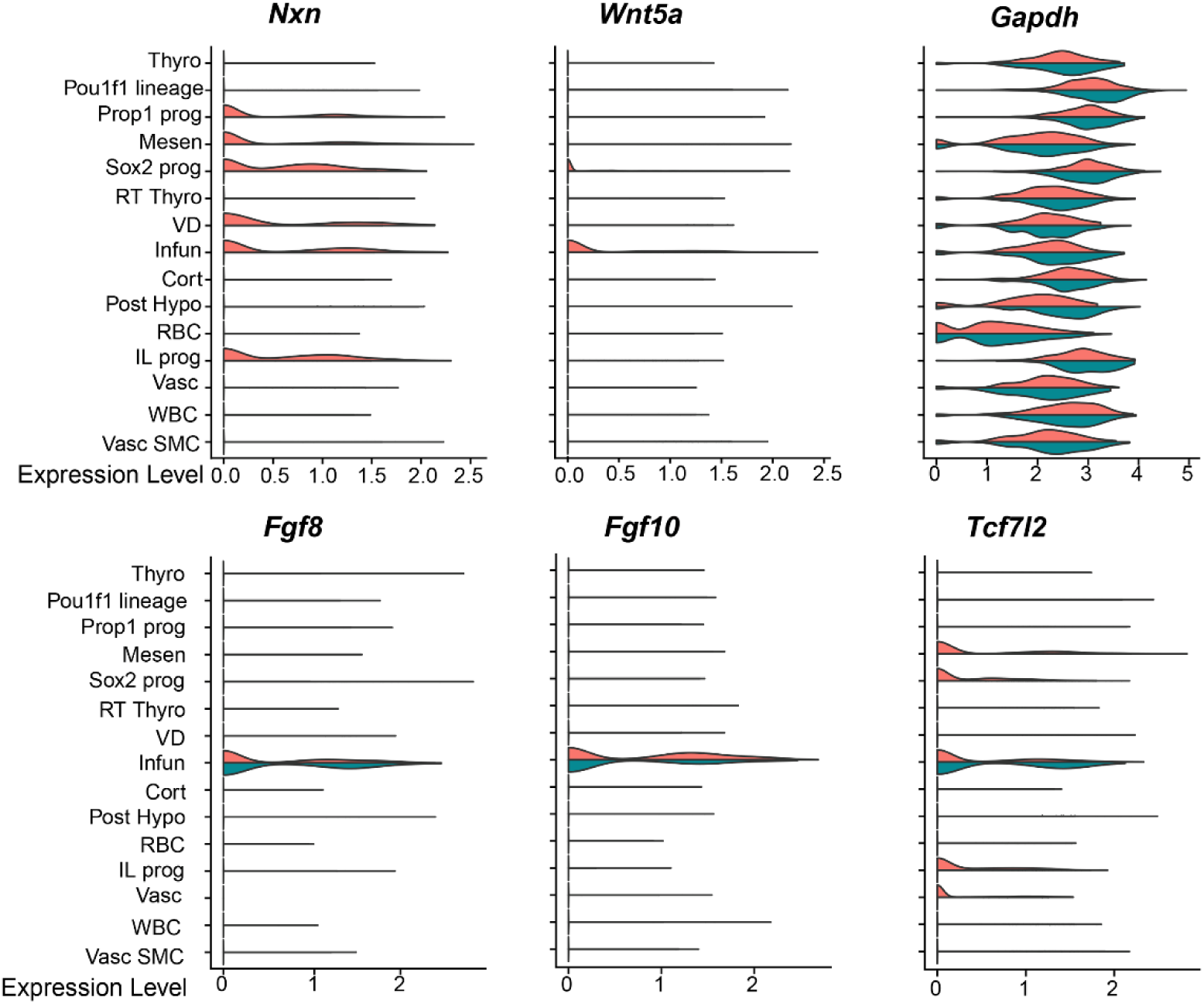
Reduced non-canonical Wnt signaling in *Nxn* mutants. Mutations in NXN or WNT5A can cause Robinow syndrome. Violin plots illustrate the detection of *Nxn* transcripts in the control scRNAseq pool (orange), but not in the mutants (green). *Nxn* transcripts are predominantly localized to the mesenchyme, neural ectoderm derived infundibulum and ventral diencephalon, and the Rathke’s pouch derived cells that include *Sox2-* expressing stem cells, and *Prop1*-expressing progenitors. Violin plots indicate a reduction in *Wnt5a* gene expression in the infundibulum of *Nxn* mutants relative to littermate controls. In contrast, the β-catenin responsive gene *Tcf7l2* is equivalently expressed in the infundibulum of *Nxn* mutants and littermate controls. There were no differences in *Fgf8* or *Fgf10* expression between *Nxn* mutants and controls, and GAPDH is expressed equivalently in both groups.

### *Nxn* deficient mice have altered WNT signaling in the infundibulum and anterior pituitary gland

Mutations in *WNT5A* are one cause of Robinow syndrome, and *Wnt5a* mutant mice have dysmorphic pituitary glands with early alterations in BMP and FGF signaling ^13,26,51^. *Wnt5a* is expressed in the infundibulum and at lower levels in the *Sox2* expressing progenitors in Rathke’s pouch (**Fig. 4**). Little or no *Wnt5a* transcripts were detected in either of these clusters in mutants. TCF7L2 function is activated by β-catenin, and *Tcf7l2* deficiency stimulates pituitary growth ^31^. *Tcf7l2* is expressed in the mesenchyme, vasculature, infundibulum, and several cell types in Rathke’s pouch, including stem cells, *Prop1* expressing progenitors, and intermediate lobe progenitors. Controls and mutants had similar levels of *Tcf7l2* transcripts in the infundibulum, but *Tcf7l2* transcripts were reduced in the other cell types in the mutants. In addition, *Nxn* deficiency did not affect *Fgf8* transcripts in the infundibulum. Thus, *Nxn* deficiency reduces *Wnt5a* expression in the infundibulum and *Tcf7l2* expression in Rathke’s pouch-derived cells.

### *Nxn* deficiency affects the proportions of Rathke’s pouch derived cell types

We re-clustered the scRNA seq data using *Epcam1* expression to focus on Rathke’s pouch-derived cell types ^60^. We detected nine distinct clusters. At e14.5 there are *Sox2* expressing stem cells within the pars distalis that will give rise to cells that express ACTH, GH, TSH, the gonadotropins LH and FSH, and PRL ^61^. At this stage intermediate lobe progenitors still express *Sox2,* and they have not yet activated detectable expression of *Pax7* or *Pomc.* However, they are distinguished from the other stem cells by higher expression of *Vax1* and *Npy* (**Fig. 5**.). We detected two clusters of progenitor cells expressing *Prop1* (a, b) and two expressing *Pou1f1* (a, b). We detected pars distalis thyrotropes expressing *Pou1f1, Cga,* and *Tshb.* We also detected the *Pou1f1-*independent rostral tip thyrotropes, and corticotropes. *Gh, Prl, Nr5a1, Lhb,* and *Fshb* transcripts were not detected at this age^62^. We compared the cell types between control and mutants and identified significant changes in the number of cells in some, but not all cell types (**Fig. 6A**). For example, the *Sox2* expressing progenitors and intermediate lobe progenitors are similar in number between the two genotypes. However, the *Prop1* a cluster, expressing *Aldh1a2* and *Ccne1,* was more abundant in mutants (28.2%) than controls (15.2%). The abundance of the *Pou1f1* a cluster was also elevated in mutants (29.4%) relative to controls (8.4%). In contrast, the clusters designated as *Prop1* b*, Pou1f1* b, pars distalis thyrotropes, and rostral tip thyrotropes were less abundant in mutants than controls [*Prop1* b, 6.5% (M), 10.7% (C); *Pou1f1* b, 2.1% (M), 19.9% (C); pars distalis thyrotropes 5.3% (M), 7.2% (C); rostral tip thyrotropes, 5.8% (M), 14.6% (C)]. Because *Nxn* is more strongly expressed in the progenitors than in more differentiated cells like rostral tip thyrotropes and *Pou1f1* expressing cells, the skewing of cell proportions in mutants relative to controls suggests that the *Nxn* deficiency is slowing the progression of progenitors to more differentiated cell fates of the rostral tip thyrotrope and *Pou1f1* lineages.

**Figure 5:**
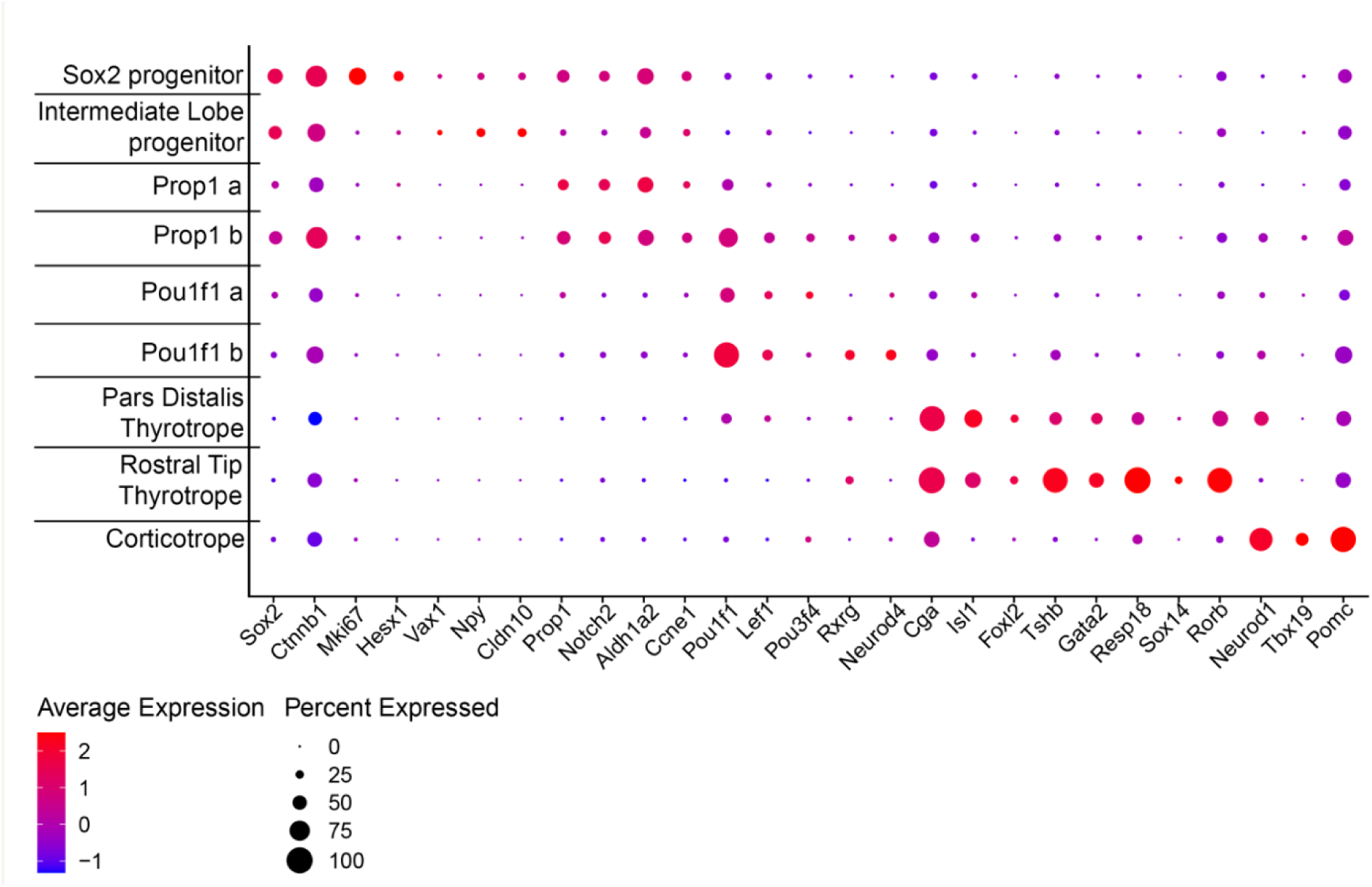
Gene expression in Rathke’s pouch derived cells defines cell clusters. Rathke’s pouch derived cells were identified based on *Epcam1* expression and re-clustered. Nine distinct clusters were detected. *Sox2* is expressed in stem cells that give rise to hormone producing cells in the anterior lobe (*Sox2* progenitors) and the intermediate lobe (Intermediate lobe progenitors). There are two clusters of cells with strong *Prop1* expression, (*Prop1 a* and *Prop1 b*). The *Prop1 b* cluster expresses higher levels of *Pou1f1* and markers of more differentiated cells like *Lef1, Rxrg,* and *Neurod4.* There are two clusters of *Pou1f1* expressing cells (*Pou1f1 a* and *Pou1f1 b*). The *Pou1f1 b* cluster has lower levels of *Prop1* than *Pou1f1 a.* The pars distalis thyrotropes and rostral tip thyrotropes (pars tuberalis) differ in that the former express *Pou1f1* and the latter express higher levels of *Sox14* and *Rorb.* Finally, corticotropes express *Neurod1, Tbx19, Pomc*.

**Figure 6.**
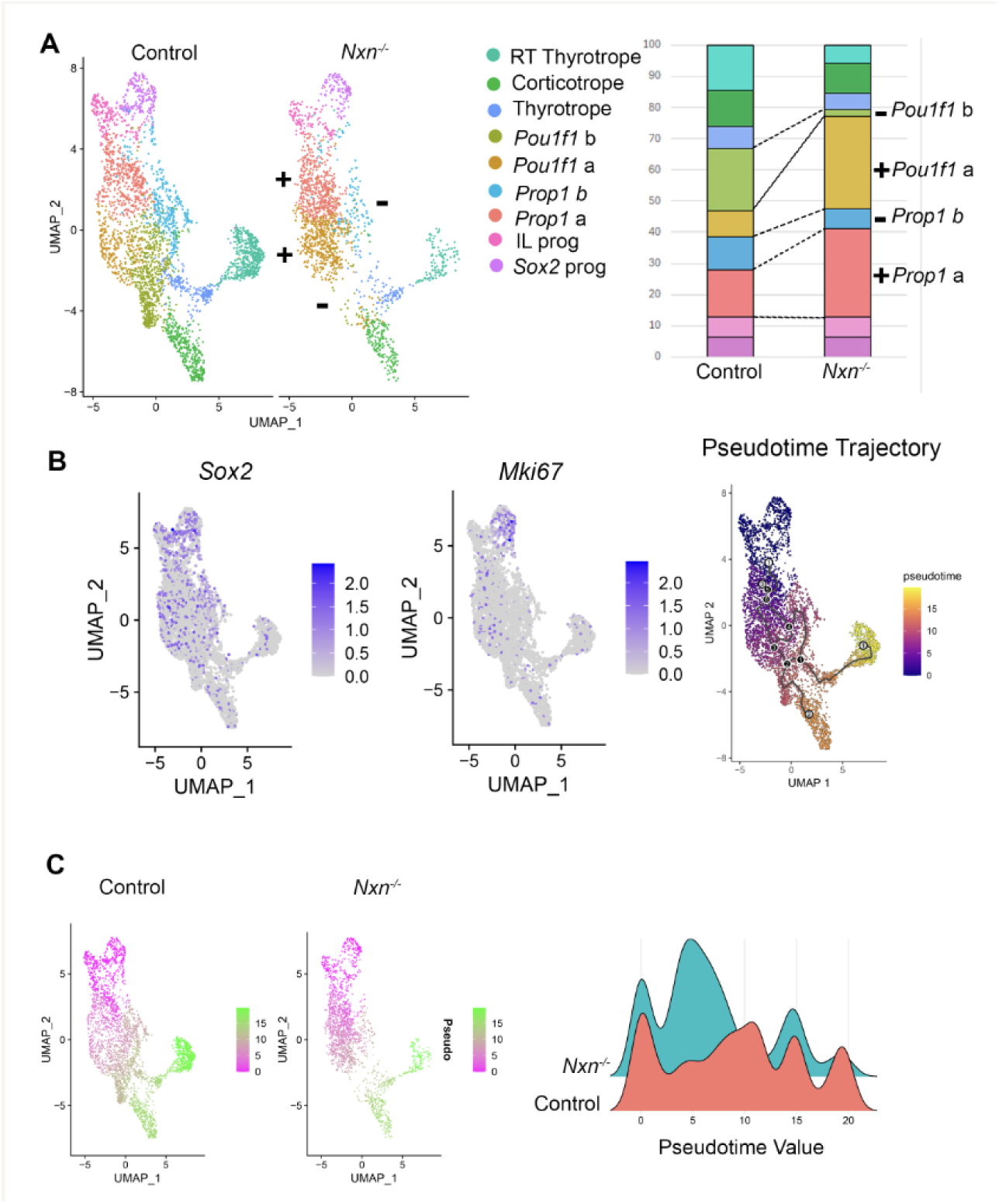
***Nxn* deficiency causes a reduced differentiation.** A. UMAP clustering of Rathke’s pouch derived cell types from e14.5 pituitaries from controls and *Nxn* mutant littermates. Stacked bar chart indicating the fraction of cells that are in each cluster. Significant differences between controls and mutants are indicated with a + (increased in mutants relative to wild type) or - (decreased). Abbreviations: rostral tip thyrotropes (RT), intermediate lobe (IL), progenitor (prog). B. UMAP plots indicating expression of the stem cell marker *Sox2* and the proliferation marker, *Mki67,* which provide the root for the pseudotime trajectory. C. Pseudotime analysis UMAP plots illustrate the progression of Rathke’s pouch-derived cell types from *Sox2*-expressing stem cells to differentiated cell types. The density plot compares distribution of cells along pseudotime axis.

We used pseudotime software to predict the trajectory of *Sox2-*expressing stem cells, which are highly proliferative at e14.5, into differentiating cells that have exited the cell cycle and exhibit reduced expression of *Mik67* (**Fig. 6B**). The pseudotime trajectory is consistent with expectations based on spatial and temporal expression of cell type specific markers. *Sox2-* expressing pituitary stem cells give rise to intermediate lobe progenitors that differentiate into *Pomc-*expressing cells later in development. The *Sox2-*expressing cells also progress to express *Prop1,* and later commit to the *Pou1f1* lineage, which is just emerging at e14.5 ^34,63^.

Some pars distalis thyrotropes are detectable by *Tshb* and *Pou1f1* transcripts at e14.5. Later in development the *Pou1f1* cells will differentiate into cells producing GH (somatotropes) and PRL (lactotropes). The first cell types to differentiate in pituitary development are the rostral tip thyrotropes, defined by expression of *Gata2* and *Tshb,* but not *Pou1f1.* These cells are readily identifiable and map at a distance from the stem cell cluster (**Fig. 6B**). Corticotropes, defined by *Tbx19* and *Pomc* expression, are also present at e14.5 and map far from the stem cell cluster.

There was a clear, genotypic difference in pseudotime values when the UMAP was split by genotype (**Fig. 6C**). We plotted pseudotime by density (scaled cell numbers), which reveals 3-4 distinct peaks. The peak representing the root, *Sox2*-expressing progenitors, is similar between genotypes, but the mutants have more cells in mid-trajectory and far fewer cells towards the end of the trajectory. Thus, pseudotime analysis reveals the paucity of more differentiated cells in mutants relative to controls. The scRNA sequencing data was also analyzed with TIME-CoExpress, a novel bioinformatic approach, which also detected a reduction in WNT signaling and a delayed differentiation of progenitor cells ^64^.

### Validation of *Nxn-*dependent changes in gene expression in pituitary tissue

To validate findings from scRNA seq analysis we examined expression of *Sox2, Pou1f1,* and *Lef1* in control and mutant tissue at e14.5 (**Fig. 7**). As expected, *Sox2* is expressed in the ventral diencephalon, infundibulum, and the marginal (stem cell) zone of the pituitary at e14.5. The marginal zone is dysmorphic and enlarged in mutants. Immunostaining generally revealed reduced POU1F1 and LEF1 staining in mutants relative to controls. All mutants had pituitary dysmorphology, but some had less profound reduction in POU1F1 and LEF1 immunostaining relative to controls. We stained for TSHB at e15.5 because the protein is not readily detected at e14.5, and we detected a reduction in the number of pars distalis thyrotropes (*Pou1f1*-dependent) in mutants (7%) compared to controls (10%), p<0.01. Thus, the changes in mRNA transcripts detected by scRNAseq e14.5 are reflected by the immunohistochemical staining in intact tissue at e14.5-e15.5.

**Figure 7.**
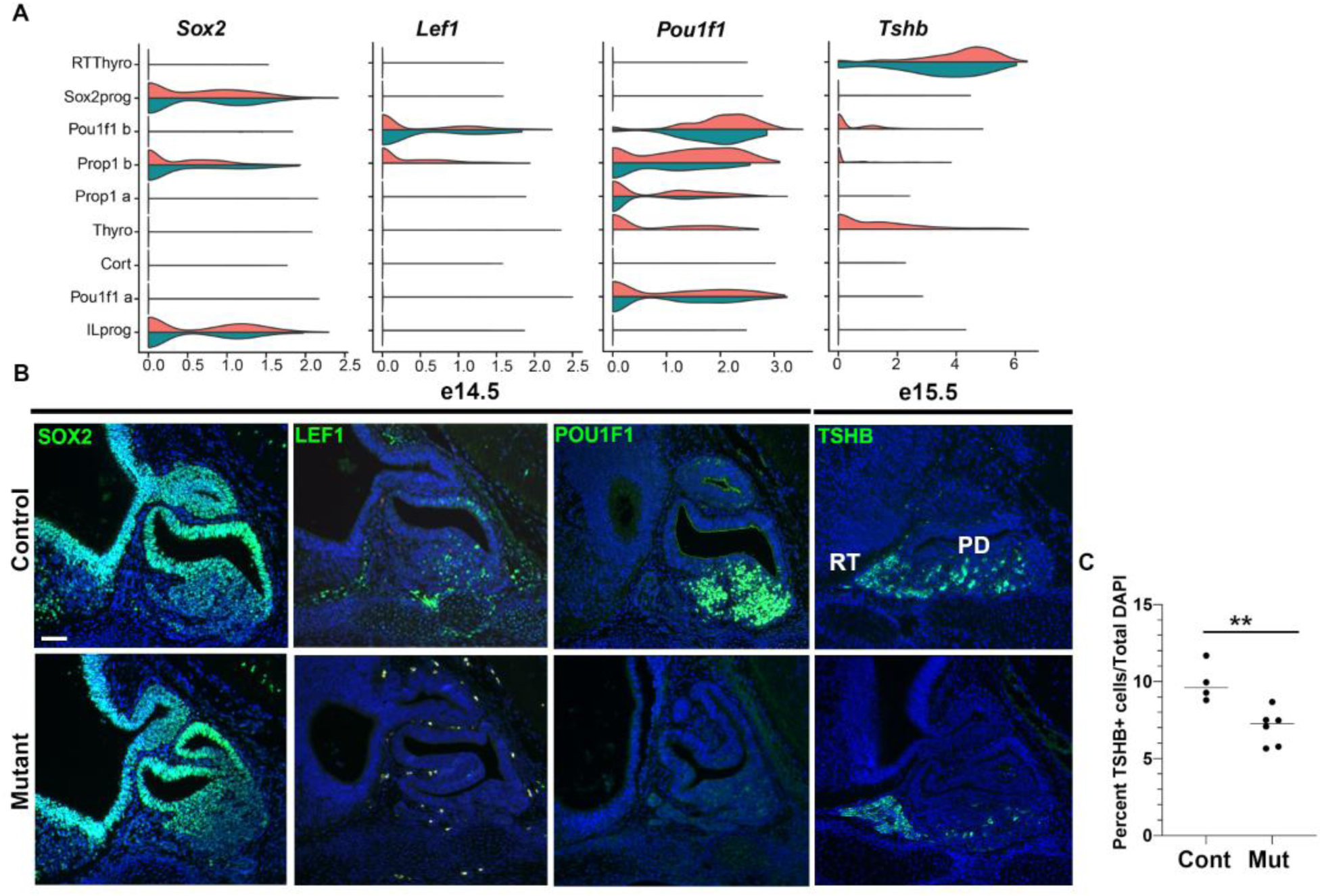
***Nxn* mutants have reduced expression of LEF1, POU1F1, and TSH, and fewer TSH producing cells.** A. Violin plots indicate expression levels of *Sox2, Lef1, Pou1f1,* and *Tshb* in controls (orange) and mutants (green). B. Sagittal sections from wild type (N=3) and *Nxn* ^-/-^ mutants (N=10) collected at e14.5 were stained with antibodies for SOX2, LEF1, and POU1F1. All mutant pituitaries exhibited dysmorphology, and 4/10 exhibited severe reduction in expression of LEF1 and POU1F1. The rostral tip thyrotropes (RT) and pars distalis thyrotropes (PD) are indicated. The scale bar= 50 uM. The p value = 0.0055 (**).

### Hormone and vasculature development appear unaffected in *Nxn* mutants

We examined pituitary development at e18.5 to determine whether the *Nxn* deficiency affected differentiation of anterior lobe hormone producing cell types that were not yet differentiated at e14.5. Histological staining revealed persistent dysmorphology of the marginal zone in mutants at e18.5 (**Supplementary Figure 4**). Immunostaining was carried out for the gonadotrope marker NR5A1, and the hormones GH, TSH and POMC. While there were no obvious differences in the pattern of immunostaining for hormones, the dysmorphic region contained cells expressing GH, TSH, and POMC. There were no obvious differences in the vasculature, as visualized by immunostaining for PECAM (**Supplementary Figure 5**).

## Conclusions

*Nxn* deficiency causes neonatal lethality, cleft palate, craniofacial abnormalities, heart defects, eye defects, and reduction in overall growth in mice ^5,52,53^ (**Supplemental Table 3**). The underlying mechanisms involve NXN-mediated effects on the pool of DVL protein ^5^. Binding of NXN to DVL prevents its interaction with other proteins, including KLHL12, a ubiquitin ligase that marks DVL for degradation ^5^. The results of *Nxn* deficiency vary by tissue. For example, the skeletal defects in *Nxn* mutants are associated with hyperactivated WNT-β-catenin signaling in osteoblasts, while the heart defects, including ventral septal defects and persistent truncus arteriosus, are associated with suppression of WNT signaling ^5^. These different effects likely result from tissue-specific differences in the pool of DVL.

We discovered pituitary dysmorphology in developing *Nxn* mutant embryos at e14.5 ^44^. The role of *Nxn* in pituitary-hypothalamic development had not been investigated. Here we report *Nxn* gene expression during pituitary development, and the effect of *Nxn* deficiency in *Nxn^em1(IMPC)J^* homozygous mice (*Nxn^-/-^*) at the earliest stages of pituitary development, including reduction in pituitary cell differentiation and reduced expression of key regulatory genes. Genes encoding members of both the canonical and non-canonical WNT signaling pathways were reduced in pituitary development. We also confirm and extend the effects of *Nxn* deficiency on other structures.

*Nxn^-/-^* exhibit embryonic lethality. Their craniofacial defects include significant shortening of head length, mandible, and nasal bone, like the *Nxn^tm1Hmik^* null homozygotes ^1^. Cleft palate, anophthalmia and microphthalmia, have been reported in *Nxn* deficient mice (**Supplemental Table 3**). The cleft palate in *Nxn^-/-^* embryos was 100% penetrant, and the majority (89%) had a complete cleft. In addition, we identified a cranial base fenestra in the basisphenoid bone. The basisphenoid bone, along with the pituitary, is a midline structure sensitive to SHH signaling ^65,66^. Inhibition of SHH signaling is sufficient to cause a fenestra in the basisphenoid bone. The basisphenoid fenestra in *Nxn^-/-^* embryos suggests that WNT signaling may also contribute to basisphenoid formation, possibly by inhibiting SHH signaling ^67^. In support of this idea, *Ptch1* and *Gli3* expression is reduced in the *Nxn* mutant pituitary progenitor cells.

*Nxn* is expressed in both the developing posterior lobe and Rathke’s pouch, and these structures were affected in *Nxn^-/-^* homozygotes. The developing posterior lobe was smaller and had reduced expression of *Wnt5a. Wnt5a* typically acts non-canonically through the WNT/Ca2+ and WNT/planar cell polarity (PCP) pathways, and it can antagonize canonical WNT signaling^68^. *Nxn* deficiency caused abnormalities in Rathke’s pouch from the onset of invagination through late gestation, especially the marginal zone where progenitors reside. This dysmorphology bears some similarity to that observed in *Wnt5a^-/-^* embryos ^13,26^. The caudal side of the pouch appears smaller in both *Nxn^-/-^* and *Wnt5a^-/-^* embryos. However, the rostral side is not expanded in *Nxn^-/-^* like *Wnt5a^-/-^* embryos. A reduction in the length of the caudal side of Rathke’s pouch is also caused by decreased SHH signaling ^65^. Because the e11.5 *Nxn^-/-^* embryos do not have the rostral expansion of pituitary progenitors, it seems unlikely that the dysmorphology observed at e12.5 and e14.5 is caused by changes in the pituitary signaling center seen in *Wnt5a^-/-^* embryos, i.e. expansion of BMP and FGF signaling ^13^. Consistent with this, we observed reduced *Wnt5a* expression in the infundibulum, but no changes in *Fgf8* or *Fgf10* expression. Thus, *Nxn* deficiency reduces non-canonical WNT signaling in the developing posterior lobe of the pituitary gland, and the planar cell polarity pathway is important for pituitary development ^69^.

The scRNA sequencing technology provides a powerful tool for phenotyping mutant embryos, and for generating hypotheses about the role of the mutant gene in normal development ^59,70^. We used scRNA sequencing to assess changes in pituitary cell populations and identify gene expression changes at e14.5 in *Nxn* mutants, and we found that *Nxn* delays pituitary progenitor cell differentiation. There are excellent markers for pituitary cell types after birth ^46,71^, but many of these are not expressed until later in gestation. We used pseudotime analysis to map the differentiation of cell types from *Sox2* expressing precursor cells. We identified cells at each stage in the developmental trajectory along with their transcriptional profile. SOX2+ stem cells transition to PROP1+ progenitor cells (*Prop1* a) that begin expression of *Pou1f1* (*Prop1* b). Each of these cell types normally express *Nxn.* The *Pou1f1* progenitors (*Pou1f1* a) have reduced *Prop1* expression and increased *Pou3f4* transcripts. They progress to a cell type the expresses *Neurod4* (*Pou1f1* b). Loss of *Nxn* delays the transition from *Prop1* a to *Prop1* b progenitors. Cells that do progress are delayed at the *Pou1f1* a progenitor stage. *Nxn^-/-^* embryos have decreased expression of *Lef1* and *Pou1f1* at the RNA and protein levels. PROP1, LEF1, and POU1F1 interact with β-catenin to regulate transcription ^32,33^; therefore, the delayed differentiation at each stage is likely caused by reduced canonical WNT signaling.

All individuals reported with Robinow syndrome and *NXN* variants appear to have biallelic loss of function alleles (**Supplemental Table 4**) ^9,51^. They all exhibit craniofacial abnormalities, skeletal dysplasia including mesomelia, and brachydactyly. They differ in presentation of cleft palate, even between siblings. A *NXN* patient homozygous for p.Arg209* had GH deficiency, but it is not clear whether the other patients were assessed for an endocrine contribution to their short stature. The delay in pituitary development that we observed in *Nxn-* deficient mice may underlie the short stature in some patients. The homozygous *Nxn* mice die at birth, which is attributable to their heart and palate defects ^5,52^. The individual homozygous for *NXN* p.Arg209* had a heart defect, as well as kidney anomalies, omphalocele, and two ventral hernias which were not reported in other patients. Thus, some babies with *NXN* biallelic loss of function alleles may not be viable. The variation in phenotypic features might be due to genetic background and environmental factors ^72^.

This study benefited from the phenotypic analysis conducted by the centers associated with the IMPC and the ready availability of cryopreserved null alleles of *Nxn* generated by the knockout mouse project. These resources enabled an in-depth phenotyping of pituitary gland organogenesis in *Nxn^-/-^* embryos, providing a plausible explanation for the endocrine deficiency observed in an individual with Robinow syndrome. Endocrine assessment of other children with Robinow syndrome could identify those who could benefit from hormone replacement therapy. Finally, analysis of other embryonic lethal strains with pituitary dysmorphology will enrich our understanding of the etiology of pituitary dysfunction in humans ^44^.

## Acknowledgements

Funding for this work was from University of South Carolina, University of Michigan, and the National Institutes of Health (R01 HD108156 to SAC and SWD, and R01HD097096 to SAC). We thank the Transgenic Animal Model and Advanced Genomics Core facilities at University of Michigan for rederiving the mice and performing single cell sequencing. We thank Drs. Prasov, Mishina, Pérez-Millán, Ellsworth, and Raetzman for useful comments.

## Supplementary Material

**Supplementary Table 1.**
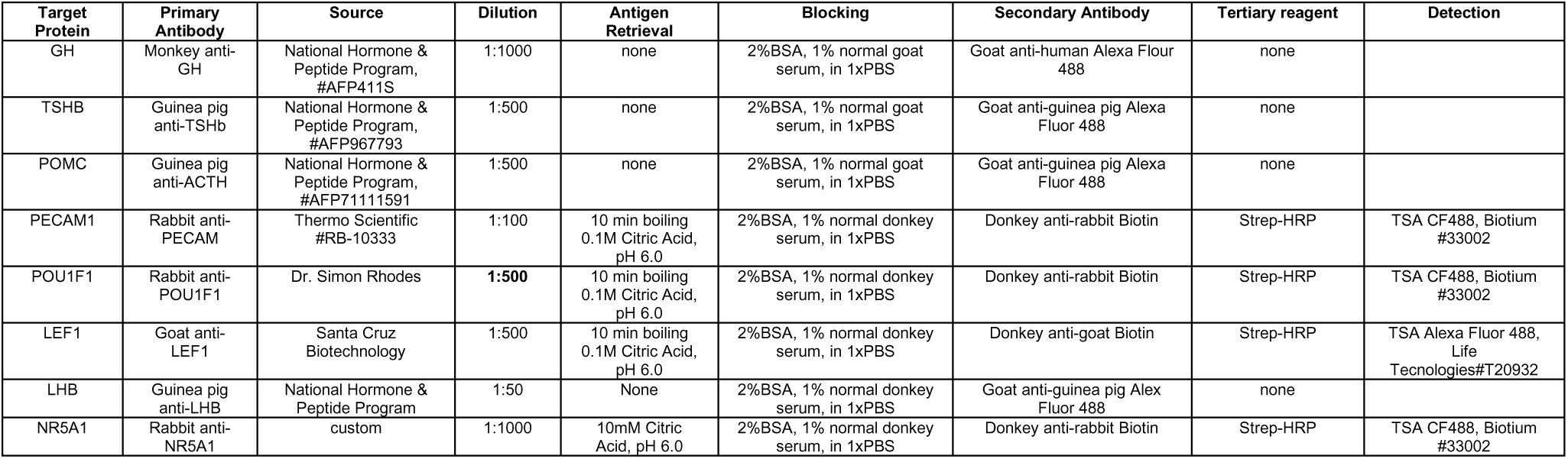
Antibody sources, dilutions and detection methods.

**Supplemental Table 2.**
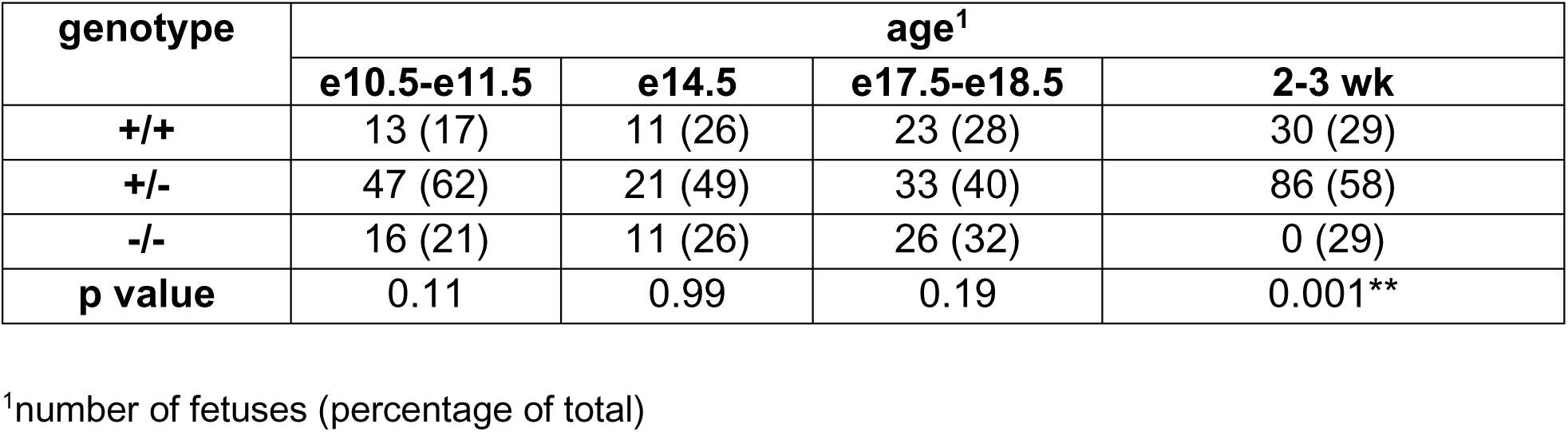
Mendelian distribution of *Nxn* genotypes during development.

**Supplemental Table 3.**
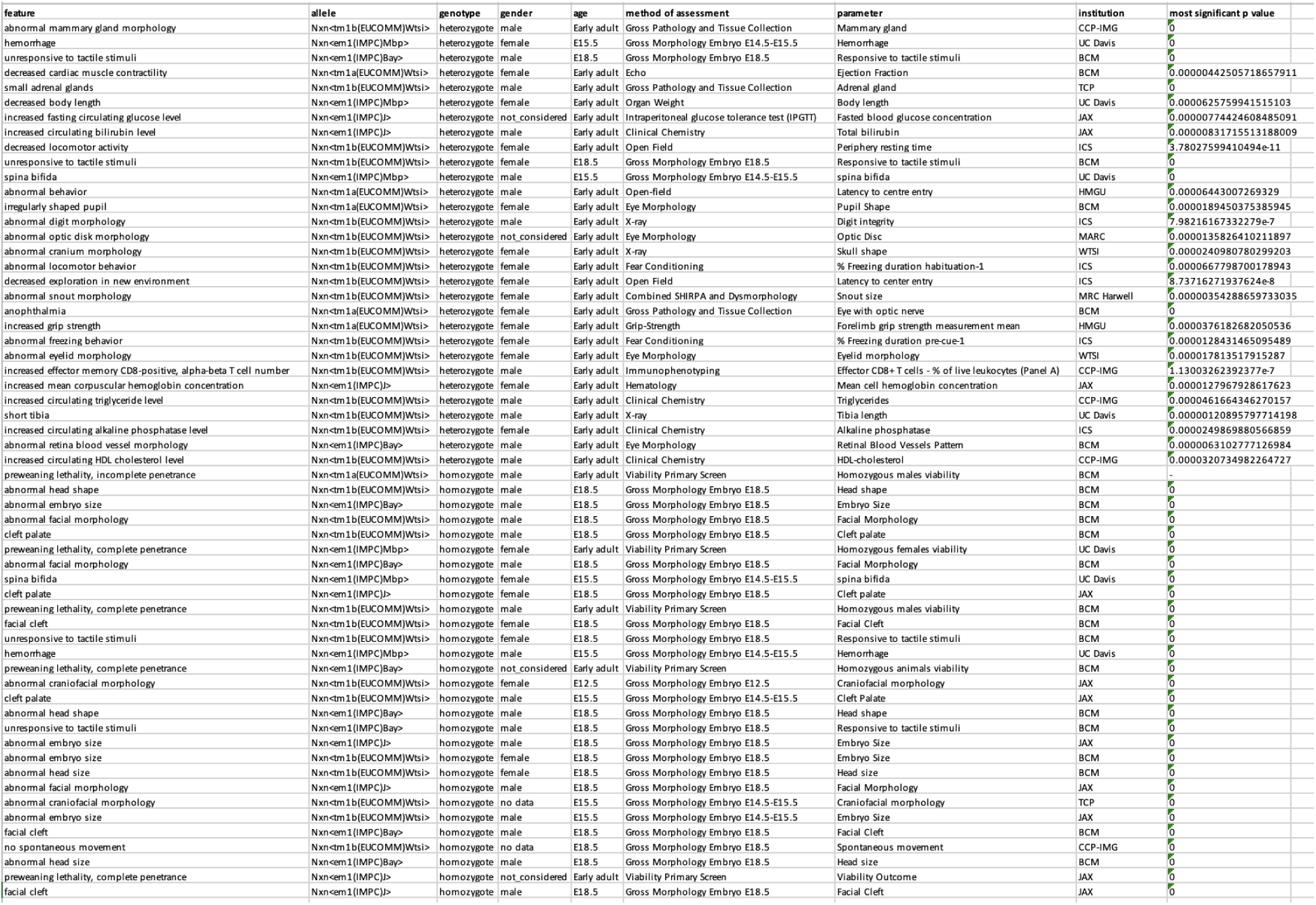
Phenotype information for homozygous and heterozygous *Nxn* alleles in mice. Multiple *Nxn* alleles have been generated and phenotyped by the International Mouse Phenotyping Consortium (IMPC) (Supplemental Fig. 1) ^73^. Phenotypes for these alleles are directly deposited with Mouse Genome Informatics (MGI) (www.informatics.jax.org/) and are available for download from the IMPC website (www.mousephenotype.org).

**Supplemental Table 4.**
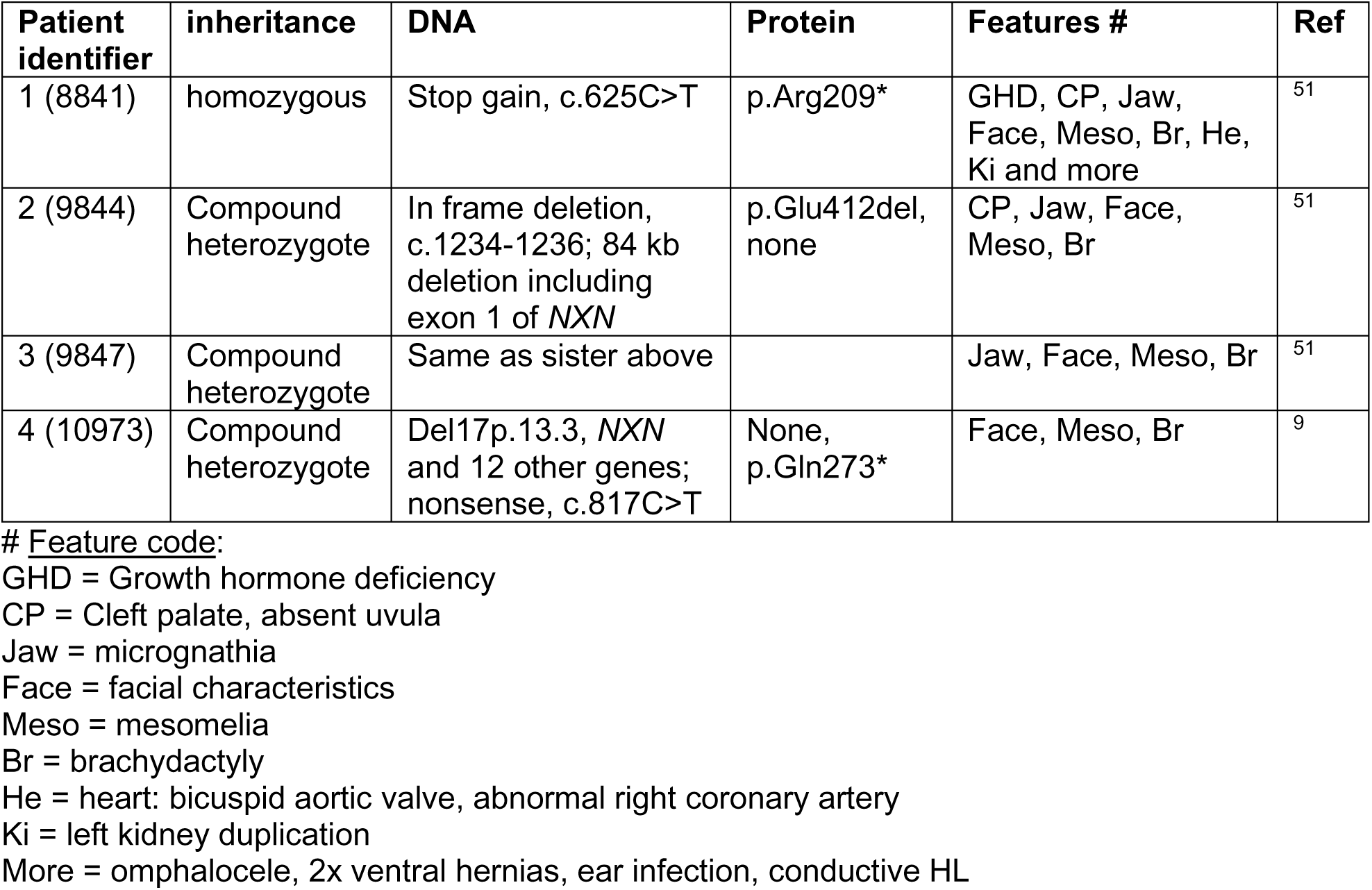
Clinical features of individuals with *NXN* variants.

**Supplemental Figure 1.**
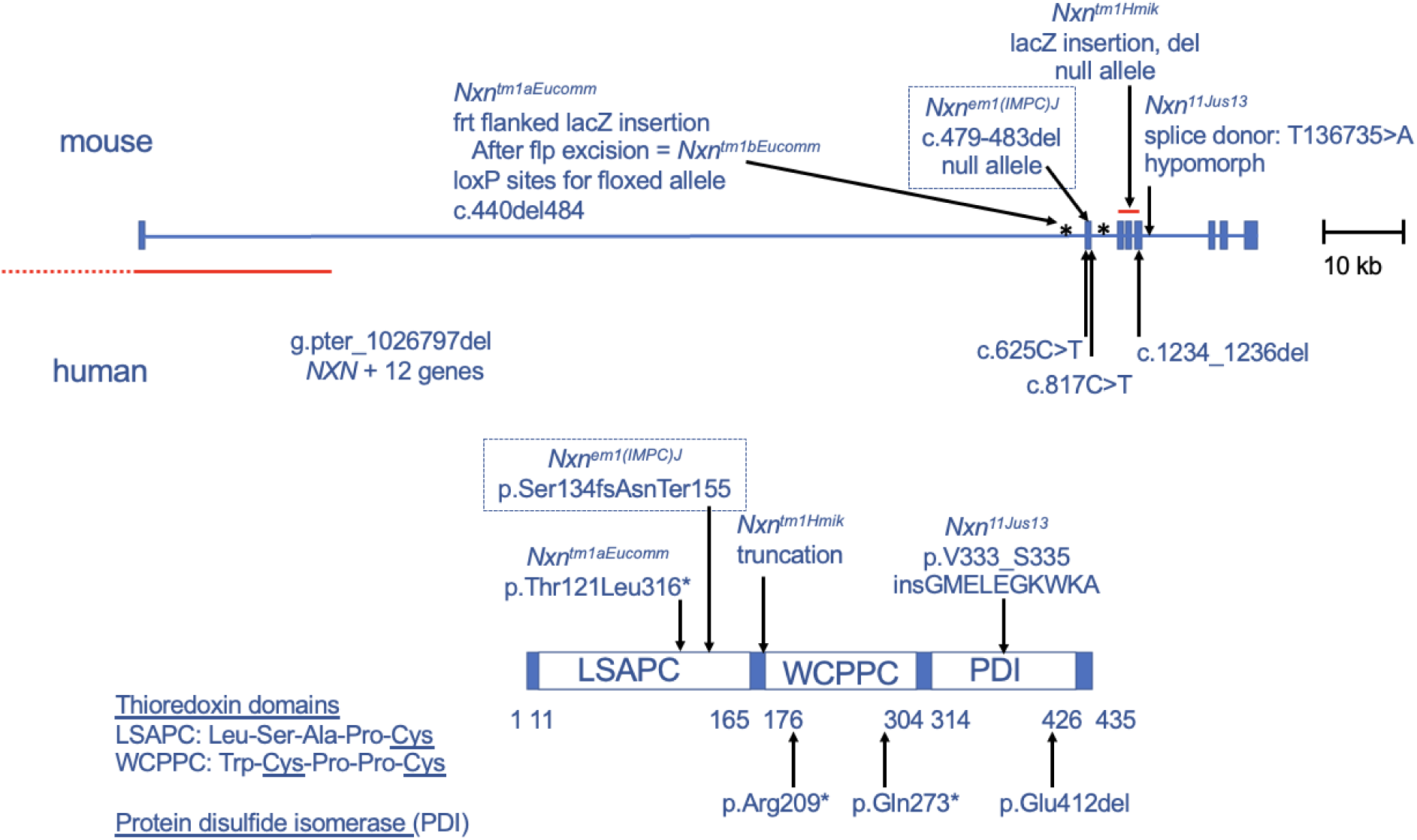
Nucleoredoxin variants in mouse and man. The structure of the human and mouse nucleoredoxin genes are similar. Mouse alleles are indicated above the gene diagram, including the knockout first Nxn^tm1aEucomm^ allele ^53^, the Nxn^em1(IMPC)J^ allele described in this manuscript, the lacZ insertion Nxn^tm1Hmik^ allele ^1^, and the hypomorphic Nxn^11Jus13^ allele ^52^. Patient mutations in NXN are indicated below the gene diagram including an individual homozygous for stop gain, c.625C>T, two compound heterozygous sisters with an 84 kb deletion including exon 1 of NXN (red) and an in frame deletion, c.1234-1236del, and an unrelated compound heterozygote with deletion of NXN and 12 other genes in trans with a nonsense mutation, c.817C>T, as described ^8,9,51^. The NXN protein consists of three domains indicated in the protein diagram. All variants in mouse and man are expected to be null alleles due to the loss of key protein domains, except the hypomorphic Nxn*^11Jus13^* allele which splices normally at a low frequency.

**Supplemental Figure 2.**
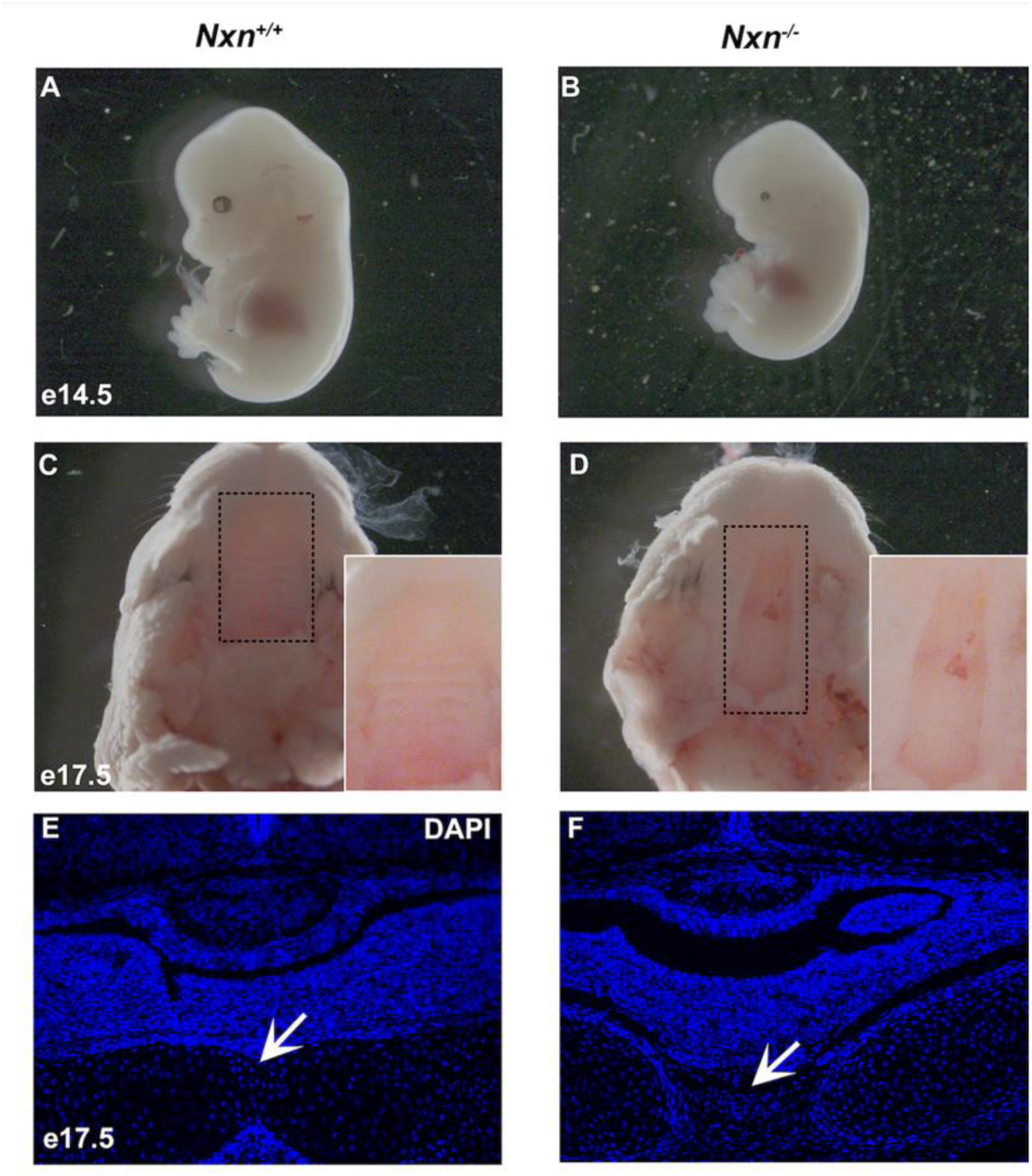
**Craniofacial abnormalities in *Nxn* mutants.** Wild type (A) and *Nxn* mutants (B) were collected at e14.5. None of the wild type animals had micro-ophthalmia or anophthalmia, but some of the mutants did. Wild type (C) and mutant (D) animals were collected at e17.5, and the palates were examined. The palate area is enlarged in the inset. Wild type (E) and mutant (F) animals were collected at e17.5, fixed, sectioned in the coronal plane, and stained with DAPI. The basisphenoid bone is fused at the base of the pituitary gland in wild type, but not mutant animals (arrows).

**Supplemental Figure 3.**
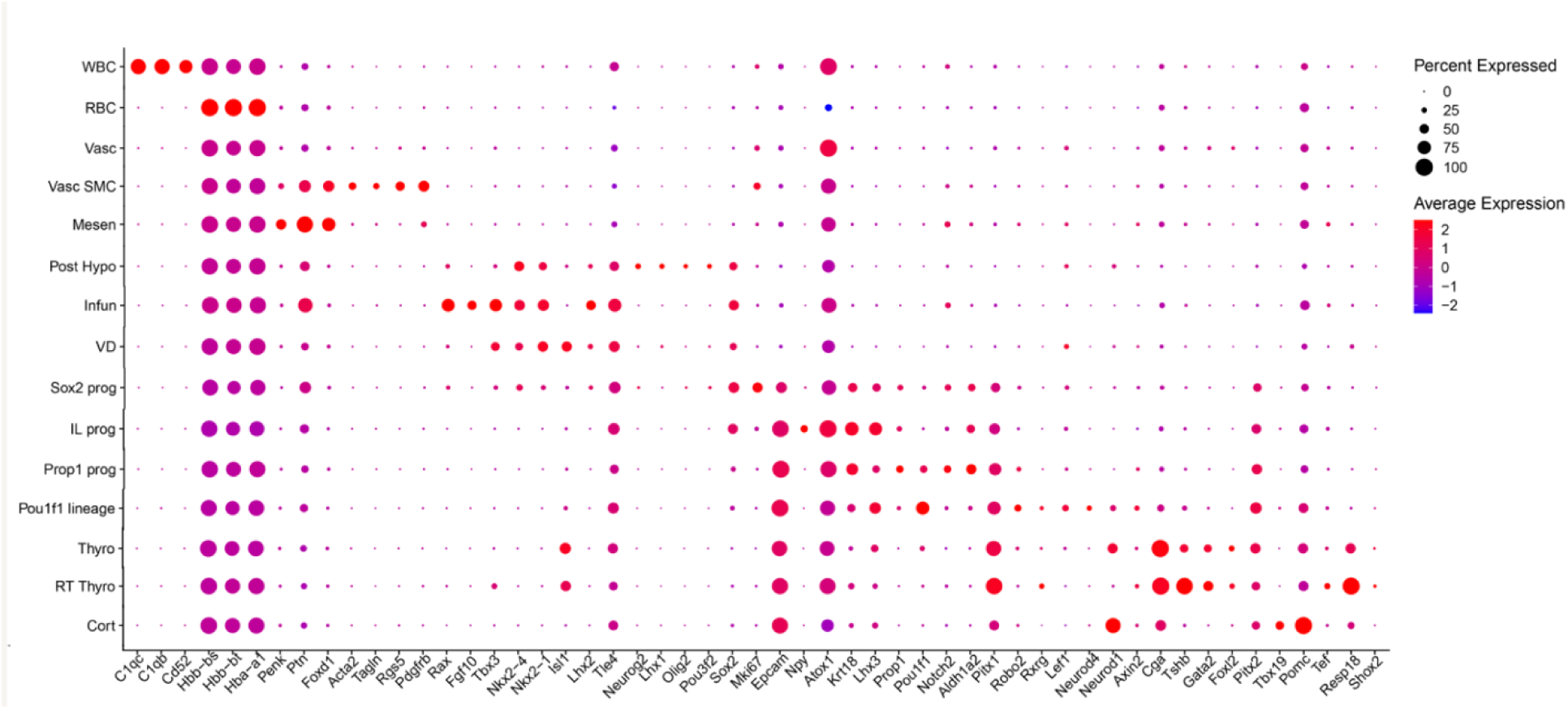
Differences in gene expression associated with cell clusters. A dot plot presents selected genes that are differentially expressed in cell clusters. Abbreviations: white blood cells (WBC), red blood cells (RBC), vasculature (Vasc), mesenchyme (Mesen), posterior hypothalamus (Post Hypo), infundibulum (Infun), ventral diencephalon (VD), progenitor (prog), intermediate lobe (IL), pars distalis thyrotrope (Thyro), rostral tip or pars tuberalis thyrotrope (RT Thyro), corticotrope (Cort).

**Supplemental Figure 4.**
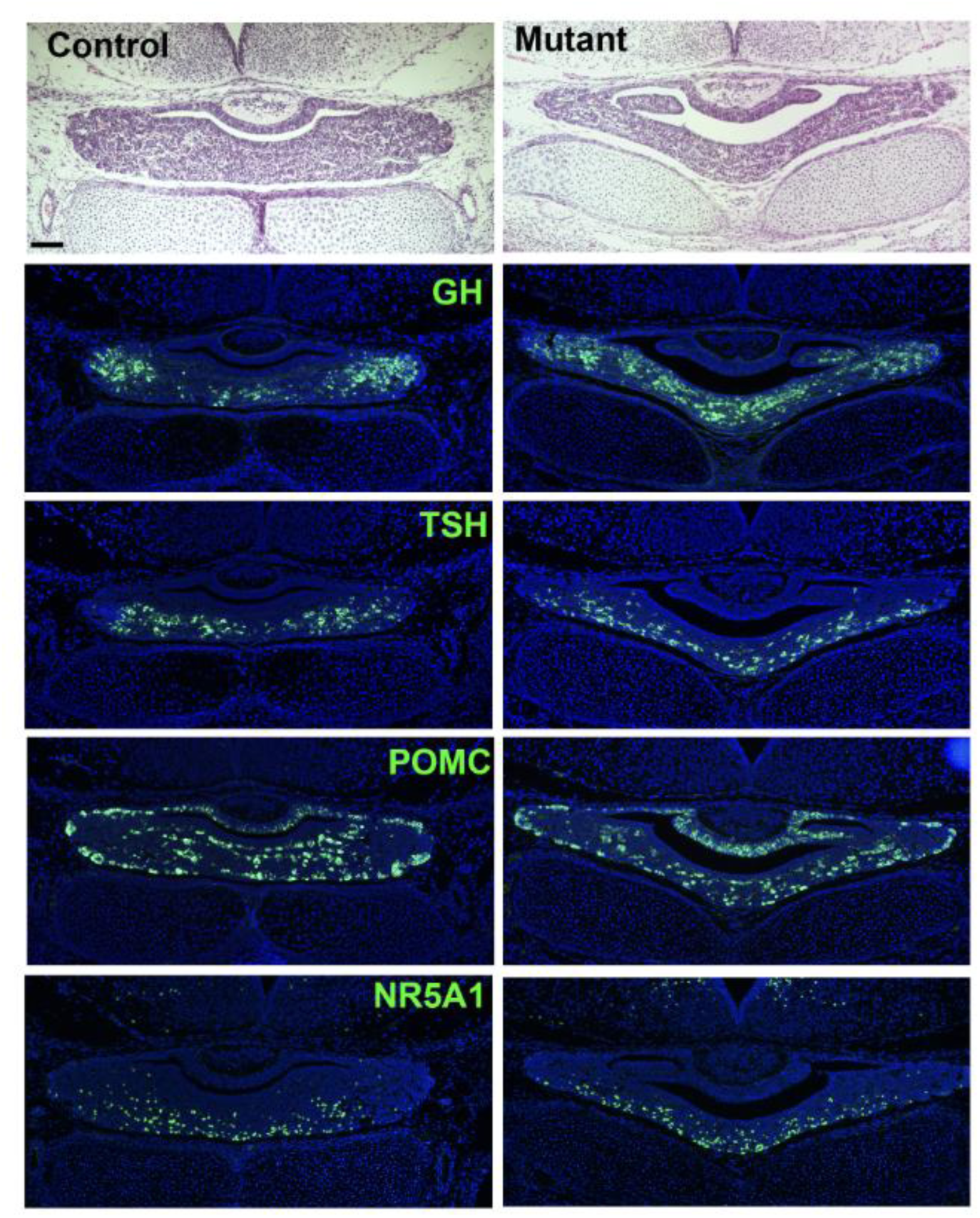
Persistent dysmorphology and differentiation of gonadotrope and *Pou1f1* lineages in later development. Pituitary glands were collected at e18.5 from controls and mutants and processed for histology by sectioning in the coronal plane. Hematoxylin and eosin staining reveals the dysmorphology that persists in the mutant pituitary. Immunostaining for NR5A1, GH, TSH and POMC did not reveal any obvious differences in mutants.

**Supplementary Figure 5.**
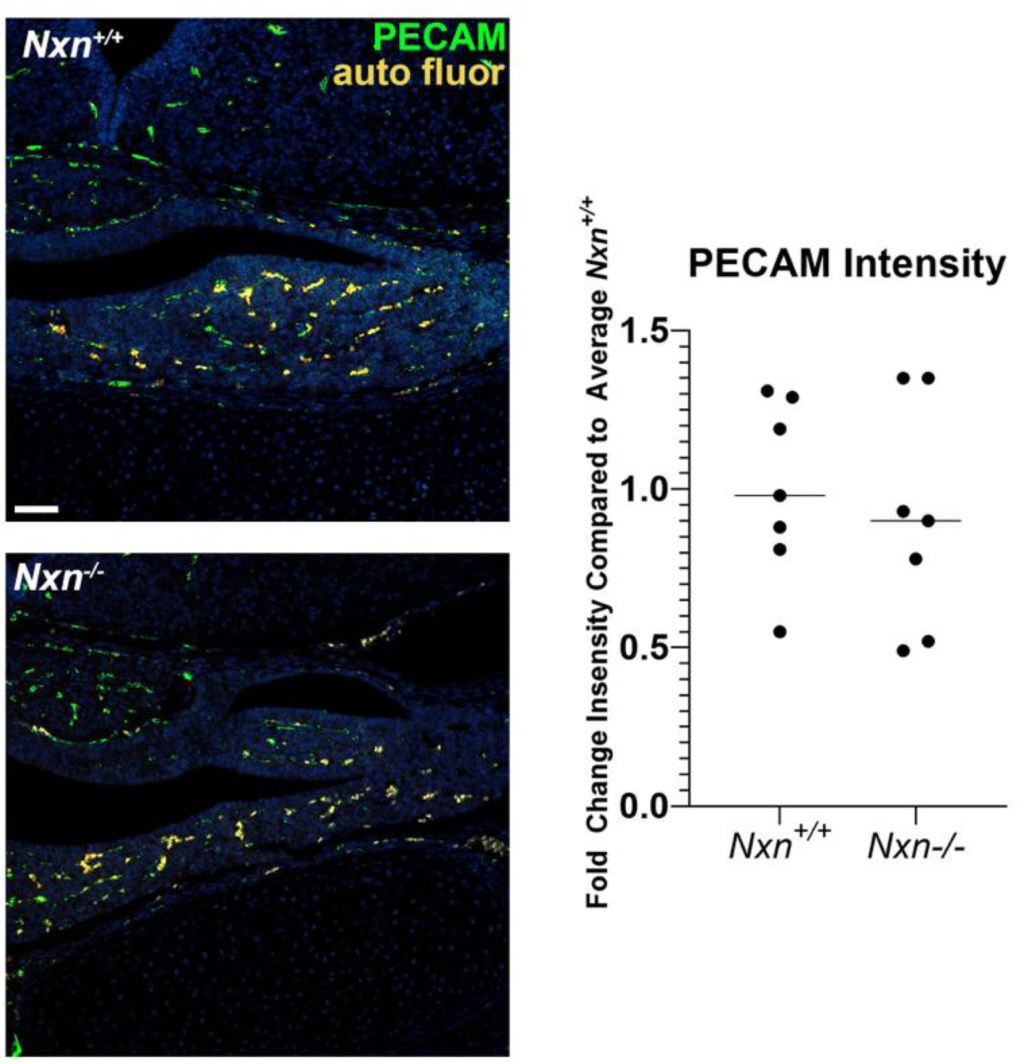
***Nxn* deficiency has no obvious effect on pituitary vasculature.** A. To assess vascular development, pituitaries were collected from e18.5 wild type fetuses and *Nxn*^-/-^ littermates, N=7/genotype. 2-5 coronal pituitary sections per individual were stained with PECAM1 antibodies (also known as CD31) and imaged by confocal microscopy at 20x with the green (Alexa488 laser) channel. Auto-fluorescent red blood cells were imaged with the red channel. Image J was used to quantify the signal intensity of the green and red channel images per area. PECAM signal intensity was calculated by subtracting the red signal from the green and averaged for the sections from a single individual. *Nxn* ^-/-^ values were normalized to the average wild type value.

