## Supplemental Material for "Nucleoredoxin regulates WNT signaling during pituitary stem cell differentiation"

### Supplementary Material

#### Supplementary Table 1.  Antibody sources, dilutions and detection methods

| Target Protein | Primary Antibody | Source | Dilution | Antigen Retrieval | Blocking | Secondary Antibody | Tertiary reagent | Detection |
|---|---|---|---|---|---|---|---|---|
| GH | Monkey anti-GH | National Hormone & Peptide Program, #AFP411S | 1:1000 | none | 2%BSA, 1% normal goat serum, in 1xPBS | Goat anti-human Alexa Flour 488 | none | |
| TSHB | Guinea pig anti-TSHb | National Hormone & Peptide Program, #AFP967793 | 1:500 | none | 2%BSA, 1% normal goat serum, in 1xPBS | Goat anti-guinea pig Alexa Fluor 488 | none | |
| POMC | Guinea pig anti-ACTH | National Hormone & Peptide Program, #AFP71111591 | 1:500 | none | 2%BSA, 1% normal goat serum, in 1xPBS | Goat anti-guinea pig Alexa Fluor 488 | none | |
| PECAM1 | Rabbit anti-PECAM | Thermo Scientific #RB-10333 | 1:100 | 10 min boiling 0.1M Citric Acid, pH 6.0 | 2%BSA, 1% normal donkey serum, in 1xPBS | Donkey anti-rabbit Biotin | Strep-HRP | TSA CF488, Biotium #33002 |
| POU1F1 | Rabbit anti-POU1F1 | Dr. Simon Rhodes | **1:500** | 10 min boiling 0.1M Citric Acid, pH 6.0 | 2%BSA, 1% normal donkey serum, in 1xPBS | Donkey anti-rabbit Biotin | Strep-HRP | TSA CF488, Biotium #33002 |
| LEF1 | Goat anti-LEF1 | Santa Cruz Biotechnology | 1:500 | 10 min boiling 0.1M Citric Acid, pH 6.0 | 2%BSA, 1% normal donkey serum, in 1xPBS | Donkey anti-goat Biotin | Strep-HRP | TSA Alexa Fluor 488, Life Tecnologies#T20932 |
| LHB | Guinea pig anti-LHB | National Hormone & Peptide Program | 1:50 | None | 2%BSA, 1% normal donkey serum, in 1xPBS | Goat anti-guinea pig Alex Fluor 488 | none | |
| NR5A1 | Rabbit anti-NR5A1 | custom | 1:1000 | 10mM Citric Acid, pH 6.0 | 2%BSA, 1% normal donkey serum, in 1xPBS | Donkey anti-rabbit Biotin | Strep-HRP | TSA CF488, Biotium #33002 |

**Supplemental Table 2. Mendelian distribution of *Nxn* genotypes during development**

| genotype | age[1] | | | |
|---|---|---|---|---|
| | e10.5-e11.5 | e14.5 | e17.5-e18.5 | 2-3 wk |
| +/+ | 13 (17) | 11 (26) | 23 (28) | 30 (29) |
| +/- | 47 (62) | 21 (49) | 33 (40) | 86 (58) |
| -/- | 16 (21) | 11 (26) | 26 (32) | 0 (29) |
| p value | 0.11 | 0.99 | 0.19 | 0.001** |

[1]number of fetuses (percentage of total)

**Supplemental Table 3. Phenotype information for homozygous and heterozygous *Nxn* alleles in mice.** Multiple *Nxn* alleles have been generated and phenotyped by the International Mouse Phenotyping Consortium (IMPC) (Supplemental Fig. 1) [73]. Phenotypes for these alleles are directly deposited with Mouse Genome Informatics (MGI) (www.informatics.jax.org/) and are available for download from the IMPC website (www.mousephenotype.org).

| feature | allele | genotype | gender | age | method of assessment | parameter | institution | most significant p value |
|---|---|---|---|---|---|---|---|---|
| abnormal mammary gland morphology | Nxn<tm1b(EUCOMM)Wtsi> | heterozygote | male | Early adult | Gross Pathology and Tissue Collection | Mammary gland | CCP-IMG | 0 |
| hemorrhage | Nxn<em1(IMPC)Mbp> | heterozygote | female | E15.5 | Gross Morphology Embryo E14.5-E15.5 | Hemorrhage | UC Davis | 0 |
| unresponsive to tactile stimuli | Nxn<em1(IMPC)Bay> | heterozygote | male | E18.5 | Gross Morphology Embryo E18.5 | Responsive to tactile stimuli | BCM | 0 |
| decreased cardiac muscle contractility | Nxn<tm1a(EUCOMM)Wtsi> | heterozygote | female | Early adult | Echo | Ejection Fraction | BCM | 0.00000442505718657911 |
| small adrenal glands | Nxn<tm1b(EUCOMM)Wtsi> | heterozygote | male | Early adult | Gross Pathology and Tissue Collection | Adrenal gland | TCP | 0 |
| decreased body length | Nxn<em1(IMPC)Mbp> | heterozygote | female | Early adult | Organ Weight | Body length | UC Davis | 0.0000625759941515103 |
| increased fasting circulating glucose level | Nxn<em1(IMPC)J> | heterozygote | not_considered | Early adult | Intraperitoneal glucose tolerance test (IPGTT) | Fasted blood glucose concentration | JAX | 0.00000774424608485091 |
| increased circulating bilirubin level | Nxn<em1(IMPC)J> | heterozygote | male | Early adult | Clinical Chemistry | Total bilirubin | JAX | 0.00000831715513188009 |
| decreased locomotor activity | Nxn<tm1b(EUCOMM)Wtsi> | heterozygote | female | Early adult | Open Field | Periphery resting time | ICS | 3.78027599410494e-11 |
| unresponsive to tactile stimuli | Nxn<tm1b(EUCOMM)Wtsi> | heterozygote | female | E18.5 | Gross Morphology Embryo E18.5 | Responsive to tactile stimuli | BCM | 0 |
| spina bifida | Nxn<em1(IMPC)Mbp> | heterozygote | male | E15.5 | Gross Morphology Embryo E14.5-E15.5 | spina bifida | UC Davis | 0 |
| abnormal behavior | Nxn<tm1a(EUCOMM)Wtsi> | heterozygote | male | Early adult | Open-field | Latency to centre entry | HMGU | 0.00006443007269329 |
| irregularly shaped pupil | Nxn<tm1a(EUCOMM)Wtsi> | heterozygote | female | Early adult | Eye Morphology | Pupil Shape | BCM | 0.0000189450375385945 |
| abnormal digit morphology | Nxn<tm1b(EUCOMM)Wtsi> | heterozygote | male | Early adult | X-ray | Digit integrity | ICS | 7.98216167332279e-7 |
| abnormal optic disk morphology | Nxn<tm1b(EUCOMM)Wtsi> | heterozygote | not_considered | Early adult | Eye Morphology | Optic Disc | MARC | 0.0000135826410211897 |
| abnormal cranium morphology | Nxn<tm1b(EUCOMM)Wtsi> | heterozygote | female | Early adult | X-ray | Skull shape | WTSI | 0.0000240980780299203 |
| abnormal locomotor behavior | Nxn<tm1b(EUCOMM)Wtsi> | heterozygote | female | Early adult | Fear Conditioning | % Freezing duration habituation-1 | ICS | 0.0000667798700178943 |
| decreased exploration in new environment | Nxn<tm1b(EUCOMM)Wtsi> | heterozygote | female | Early adult | Open Field | Latency to center entry | ICS | 8.73716271937624e-8 |
| abnormal snout morphology | Nxn<tm1b(EUCOMM)Wtsi> | heterozygote | female | Early adult | Combined SHIRPA and Dysmorphology | Snout size | MRC Harwell | 0.00000354288659733035 |
| anophthalmia | Nxn<tm1a(EUCOMM)Wtsi> | heterozygote | female | Early adult | Gross Pathology and Tissue Collection | Eye with optic nerve | BCM | 0 |
| increased grip strength | Nxn<tm1a(EUCOMM)Wtsi> | heterozygote | female | Early adult | Grip-Strength | Forelimb grip strength measurement mean | HMGU | 0.0000376182682050536 |
| abnormal freezing behavior | Nxn<tm1b(EUCOMM)Wtsi> | heterozygote | female | Early adult | Fear Conditioning | % Freezing duration pre-cue-1 | ICS | 0.0000128431465095489 |
| abnormal eyelid morphology | Nxn<tm1b(EUCOMM)Wtsi> | heterozygote | female | Early adult | Eye Morphology | Eyelid morphology | WTSI | 0.000017813517915287 |
| increased effector memory CD8-positive, alpha-beta T cell number | Nxn<tm1b(EUCOMM)Wtsi> | heterozygote | male | Early adult | Immunophenotyping | Effector CD8+ T cells - % of live leukocytes (Panel A) | CCP-IMG | 1.13003262392377e-7 |
| increased mean corpuscular hemoglobin concentration | Nxn<em1(IMPC)J> | heterozygote | female | Early adult | Hematology | Mean cell hemoglobin concentration | JAX | 0.000127967928617623 |
| increased circulating triglyceride level | Nxn<tm1b(EUCOMM)Wtsi> | heterozygote | male | Early adult | Clinical Chemistry | Triglycerides | CCP-IMG | 0.0000461664346270157 |
| short tibia | Nxn<tm1b(EUCOMM)Wtsi> | heterozygote | male | Early adult | X-ray | Tibia length | UC Davis | 0.00000120895797714198 |
| increased circulating alkaline phosphatase level | Nxn<tm1b(EUCOMM)Wtsi> | heterozygote | female | Early adult | Clinical Chemistry | Alkaline phosphatase | ICS | 0.0000249869880566859 |
| abnormal retina blood vessel morphology | Nxn<em1(IMPC)Bay> | heterozygote | male | Early adult | Eye Morphology | Retinal Blood Vessels Pattern | BCM | 0.0000063102777126984 |
| increased circulating HDL cholesterol level | Nxn<tm1b(EUCOMM)Wtsi> | heterozygote | male | Early adult | Clinical Chemistry | HDL-cholesterol | CCP-IMG | 0.0000320734982264727 |
| preweaning lethality, incomplete penetrance | Nxn<tm1a(EUCOMM)Wtsi> | homozygote | male | Early adult | Viability Primary Screen | Homozygous males viability | BCM | - |
| abnormal head shape | Nxn<tm1b(EUCOMM)Wtsi> | homozygote | male | E18.5 | Gross Morphology Embryo E18.5 | Head shape | BCM | 0 |
| abnormal embryo size | Nxn<em1(IMPC)Bay> | homozygote | male | E18.5 | Gross Morphology Embryo E18.5 | Embryo Size | BCM | 0 |
| abnormal facial morphology | Nxn<tm1b(EUCOMM)Wtsi> | homozygote | male | E18.5 | Gross Morphology Embryo E18.5 | Facial Morphology | BCM | 0 |
| cleft palate | Nxn<tm1b(EUCOMM)Wtsi> | homozygote | male | E18.5 | Gross Morphology Embryo E18.5 | Cleft palate | BCM | 0 |
| preweaning lethality, complete penetrance | Nxn<em1(IMPC)Mbp> | homozygote | female | Early adult | Viability Primary Screen | Homozygous females viability | UC Davis | 0 |
| abnormal facial morphology | Nxn<em1(IMPC)Bay> | homozygote | male | E18.5 | Gross Morphology Embryo E18.5 | Facial Morphology | BCM | 0 |
| spina bifida | Nxn<em1(IMPC)Mbp> | homozygote | female | E15.5 | Gross Morphology Embryo E14.5-E15.5 | spina bifida | UC Davis | 0 |
| cleft palate | Nxn<em1(IMPC)J> | homozygote | female | E18.5 | Gross Morphology Embryo E18.5 | Cleft palate | JAX | 0 |
| preweaning lethality, complete penetrance | Nxn<tm1b(EUCOMM)Wtsi> | homozygote | male | Early adult | Viability Primary Screen | Homozygous males viability | BCM | 0 |
| facial cleft | Nxn<tm1b(EUCOMM)Wtsi> | homozygote | female | E18.5 | Gross Morphology Embryo E18.5 | Facial Cleft | BCM | 0 |
| unresponsive to tactile stimuli | Nxn<tm1b(EUCOMM)Wtsi> | homozygote | female | E18.5 | Gross Morphology Embryo E18.5 | Responsive to tactile stimuli | BCM | 0 |
| hemorrhage | Nxn<em1(IMPC)Mbp> | homozygote | male | E15.5 | Gross Morphology Embryo E14.5-E15.5 | Hemorrhage | UC Davis | 0 |
| preweaning lethality, complete penetrance | Nxn<em1(IMPC)Bay> | homozygote | not_considered | Early adult | Viability Primary Screen | Homozygous animals viability | BCM | 0 |
| abnormal craniofacial morphology | Nxn<tm1b(EUCOMM)Wtsi> | homozygote | female | E12.5 | Gross Morphology Embryo E12.5 | Craniofacial morphology | JAX | 0 |
| cleft palate | Nxn<tm1b(EUCOMM)Wtsi> | homozygote | male | E15.5 | Gross Morphology Embryo E14.5-E15.5 | Cleft Palate | JAX | 0 |
| abnormal head shape | Nxn<em1(IMPC)Bay> | homozygote | male | E18.5 | Gross Morphology Embryo E18.5 | Head shape | BCM | 0 |
| unresponsive to tactile stimuli | Nxn<em1(IMPC)Bay> | homozygote | male | E18.5 | Gross Morphology Embryo E18.5 | Responsive to tactile stimuli | BCM | 0 |
| abnormal embryo size | Nxn<em1(IMPC)J> | homozygote | male | E18.5 | Gross Morphology Embryo E18.5 | Embryo Size | JAX | 0 |
| abnormal embryo size | Nxn<tm1b(EUCOMM)Wtsi> | homozygote | female | E18.5 | Gross Morphology Embryo E18.5 | Embryo Size | BCM | 0 |
| abnormal head size | Nxn<tm1b(EUCOMM)Wtsi> | homozygote | female | E18.5 | Gross Morphology Embryo E18.5 | Head size | BCM | 0 |
| abnormal facial morphology | Nxn<em1(IMPC)J> | homozygote | male | E18.5 | Gross Morphology Embryo E18.5 | Facial Morphology | JAX | 0 |
| abnormal craniofacial morphology | Nxn<tm1b(EUCOMM)Wtsi> | homozygote | no data | E15.5 | Gross Morphology Embryo E14.5-E15.5 | Craniofacial morphology | TCP | 0 |
| abnormal embryo size | Nxn<tm1b(EUCOMM)Wtsi> | homozygote | male | E15.5 | Gross Morphology Embryo E14.5-E15.5 | Embryo Size | JAX | 0 |
| facial cleft | Nxn<em1(IMPC)Bay> | homozygote | male | E18.5 | Gross Morphology Embryo E18.5 | Facial Cleft | BCM | 0 |
| no spontaneous movement | Nxn<tm1b(EUCOMM)Wtsi> | homozygote | no data | E18.5 | Gross Morphology Embryo E18.5 | Spontaneous movement | CCP-IMG | 0 |
| abnormal head size | Nxn<em1(IMPC)Bay> | homozygote | male | E18.5 | Gross Morphology Embryo E18.5 | Head size | BCM | 0 |
| preweaning lethality, complete penetrance | Nxn<em1(IMPC)J> | homozygote | not_considered | Early adult | Viability Primary Screen | Viability Outcome | JAX | 0 |
| facial cleft | Nxn<em1(IMPC)J> | homozygote | male | E18.5 | Gross Morphology Embryo E18.5 | Facial Cleft | JAX | 0 |

**Supplemental Table 4. Clinical features of individuals with *NXN* variants.**

| Patient identifier | inheritance | DNA | Protein | Features # | Ref |
|---|---|---|---|---|---|
| 1 (8841) | homozygous | Stop gain, c.625C>T | p.Arg209* | GHD, CP, Jaw, Face, Meso, Br, He, Ki and more | [51] |
| 2 (9844) | Compound heterozygote | In frame deletion, c.1234-1236; 84 kb deletion including exon 1 of *NXN* | p.Glu412del, none | CP, Jaw, Face, Meso, Br | [51] |
| 3 (9847) | Compound heterozygote | Same as sister above | | Jaw, Face, Meso, Br | [51] |
| 4 (10973) | Compound heterozygote | Del17p.13.3, *NXN* and 12 other genes; nonsense, c.817C>T | None, p.Gln273* | Face, Meso, Br | [9] |

### Feature code:
GHD = Growth hormone deficiency
CP = Cleft palate, absent uvula
Jaw = micrognathia
Face = facial characteristics
Meso = mesomelia
Br = brachydactyly
He = heart: bicuspid aortic valve, abnormal right coronary artery
Ki = left kidney duplication
More = omphalocele, 2x ventral hernias, ear infection, conductive HL

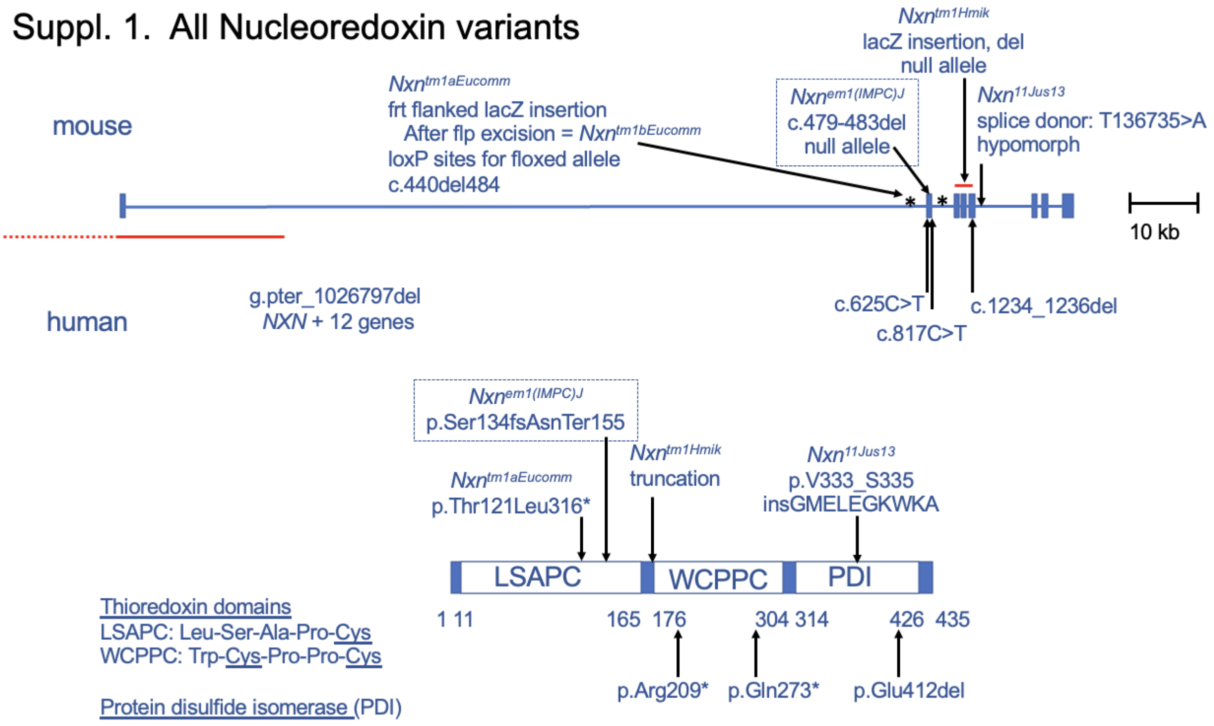


Supplemental Figure 1. Nucleoredoxin variants in mouse and man. The structure of the human and mouse nucleoredoxin genes are similar. Mouse alleles are indicated above the gene diagram, including the knockout first *Nxn^tm1aEucomm^* allele [53], the *Nxn^em1(IMPC)J^* allele described in this manuscript, the lacZ insertion *Nxn^tm1Hmik^* allele [1], and the hypomorphic *Nxn^11Jus13^* allele [52]. Patient mutations in *NXN* are indicated below the gene diagram including an individual homozygous for stop gain, c.625C>T, two compound heterozygous sisters with an 84 kb deletion including exon 1 of *NXN* (red) and an in frame deletion, c.1234-1236del, and an unrelated compound heterozygote with deletion of *NXN* and 12 other genes in trans with a nonsense mutation, c.817C>T, as described [8,9,51]. The NXN protein consists of three domains indicated in the protein diagram. All variants in mouse and man are expected to be null alleles due to the loss of key protein domains, except the hypomorphic *Nxn^11Jus13^* allele which splices normally at a low frequency.

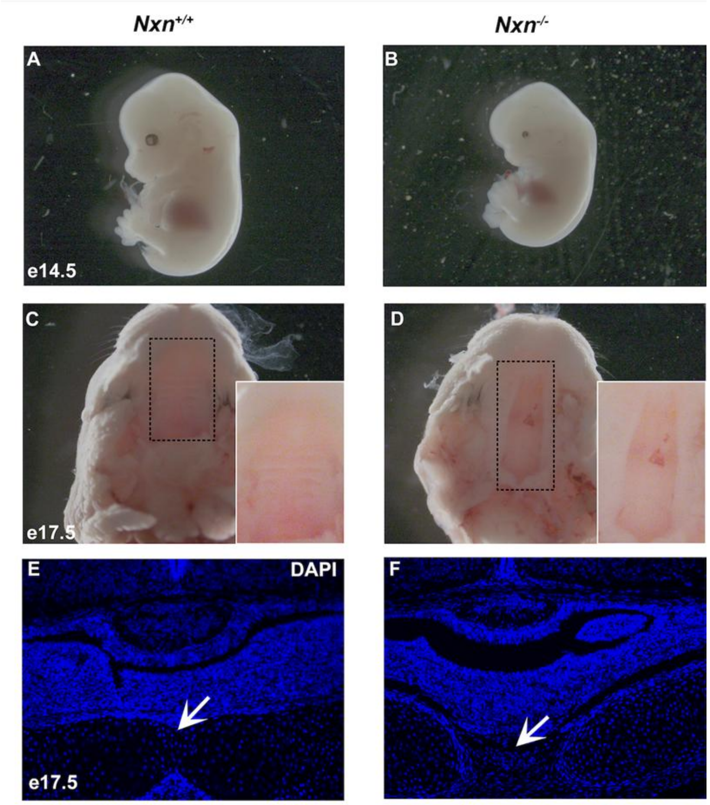


Supplemental Figure 2. Craniofacial abnormalities in *Nxn* mutants.

Wild type (A) and *Nxn* mutants (B) were collected at e14.5. None of the wild type animals had micro-ophthalmia or anophthalmia, but some of the mutants did.

Wild type (C) and mutant (D) animals were collected at e17.5, and the palates were examined. The palate area is enlarged in the inset.

Wild type (E) and mutant (F) animals were collected at e17.5, fixed, sectioned in the coronal plane, and stained with DAPI. The basisphenoid bone is fused at the base of the pituitary gland in wild type, but not mutant animals (arrows).

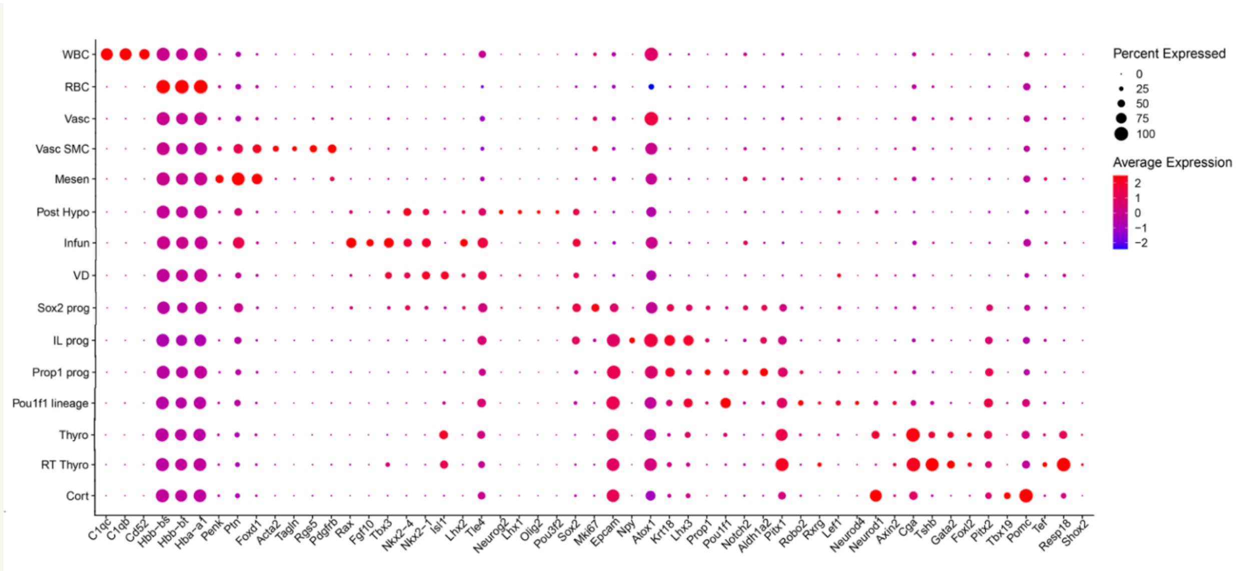


Supplemental Figure 3. Differences in gene expression associated with cell clusters.

A dot plot presents selected genes that are differentially expressed in cell clusters. Abbreviations: white blood cells (WBC), red blood cells (RBC), vasculature (Vasc), mesenchyme (Mesen), posterior hypothalamus (Post Hypo), infundibulum (Infun), ventral diencephalon (VD), progenitor (prog), intermediate lobe (IL), pars distalis thyrotrope (Thyro), rostral tip or pars tuberalis thyrotrope (RT Thyro), corticotrope (Cort).

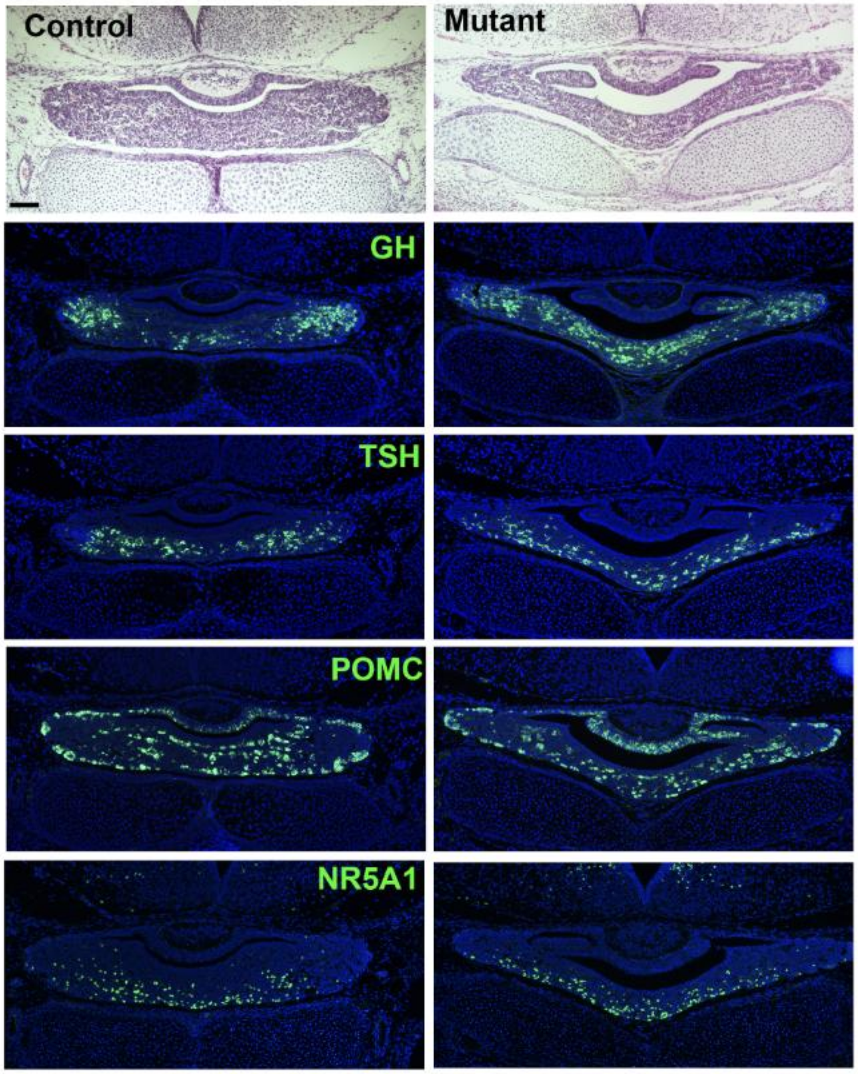


Supplemental Figure 4. Persistent dysmorphology and differentiation of gonadotrope and *Pou1f1* lineages in later development.

Pituitary glands were collected at e18.5 from controls and mutants and processed for histology by sectioning in the coronal plane. Hematoxylin and eosin staining reveals the dysmorphology that persists in the mutant pituitary. Immunostaining for NR5A1, GH, TSH and POMC did not reveal any obvious differences in mutants.

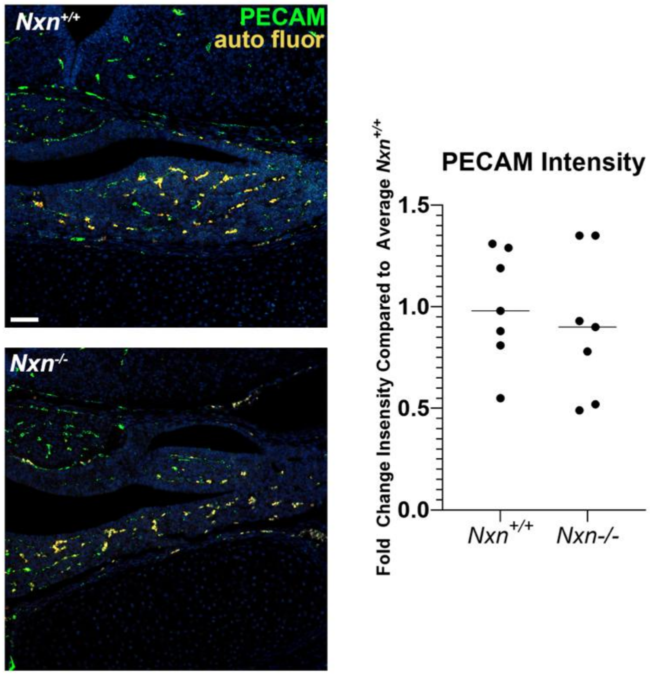


<u>Supplementary Figure 5. *Nxn* deficiency has no obvious effect on pituitary vasculature.</u>

A. To assess vascular development, pituitaries were collected from e18.5 wild type fetuses and *Nxn*$^{-/-}$ littermates, N=7/genotype. 2-5 coronal pituitary sections per individual were stained with PECAM1 antibodies (also known as CD31) and imaged by confocal microscopy at 20x with the green (Alexa488 laser) channel. Auto-fluorescent red blood cells were imaged with the red channel. Image J was used to quantify the signal intensity of the green and red channel images per area. PECAM signal intensity was calculated by subtracting the red signal from the green and averaged for the sections from a single individual. *Nxn*$^{-/-}$ values were normalized to the average wild type value.
